# Emergent 3D genome reorganization from the stepwise assembly of transcriptional condensates

**DOI:** 10.1101/2025.02.23.639564

**Authors:** Surabhi Chowdhary, Sarah Paracha, Lucas Dyer, David Pincus

## Abstract

Transcriptional condensates are clusters of transcription factors, coactivators, and RNA Pol II associated with high-level gene expression, yet how they assemble and function within the cell remains unclear. Here we show that transcriptional condensates form in a stepwise manner to enable both graded and three-dimensional (3D) gene control in the yeast heat shock response. Molecular dissection revealed a condensate cascade. First, the transcription factor Hsf1 clusters upon partial dissociation from the chaperone Hsp70. Next, the coactivator Mediator partitions following further Hsp70 dissociation and Hsf1 phosphorylation. Finally, Pol II condenses, driving the emergent coalescence of HSR genes. Molecular analysis of a series of Hsf1 mutants revealed graded, rather than switch-like, transcriptional activity. Separation-of-function mutants showed that condensate formation can be decoupled from gene activation. Instead, fully assembled HSR condensates promote adaptive 3D genome reconfiguration, suggesting a role of transcriptional condensates beyond gene activation.

## INTRODUCTION

Gene regulation is a fundamental process that underpins growth and development, and disrupted gene control is the basis for most human diseases. 3D genome organization and biomolecular condensates are key determinants of eukaryotic gene regulation, and mutations in either are causally linked to cancer, neurodegeneration, and developmental disorders ^1–3^. Biomolecular condensates are self-organizing, membrane-free macromolecular assemblies that selectively partition specific biomolecules ^4^. Compartmentalization of transcriptional machinery in biomolecular condensates at specific genomic loci is associated with robust expression of cell identity, proliferation, development, and stress-induced genes ^5–12^.

Transcriptional condensates are hypothesized to involve cooperative association of sequence-specific transcription factors, coactivators, and RNA Polymerase II (Pol II) ^5–7,9,13–17^, yet how they are assembled is unclear. There are two plausible models for how transcriptional condensates may form, each with potentially distinct functional implications (Figure 1A). The first is the concerted recruitment model, wherein constituent biomolecules are recruited in a non-stoichiometric and synergistic manner, and a network of multivalent interactions among the various components reinforces the spontaneous assembly of the condensate (Figure 1A). These low-affinity, multivalent interactions among domains with compatible surface properties are commonly attributed to intrinsically disordered regions (IDRs), which are frequently present in the constituent proteins ^18–20^. Also embedded within this network of disordered interactions are allosteric regulatory interactions between the structured domains of the proteins, which may proscribe complex kinetic control ^21–24^.

**Figure 1.**
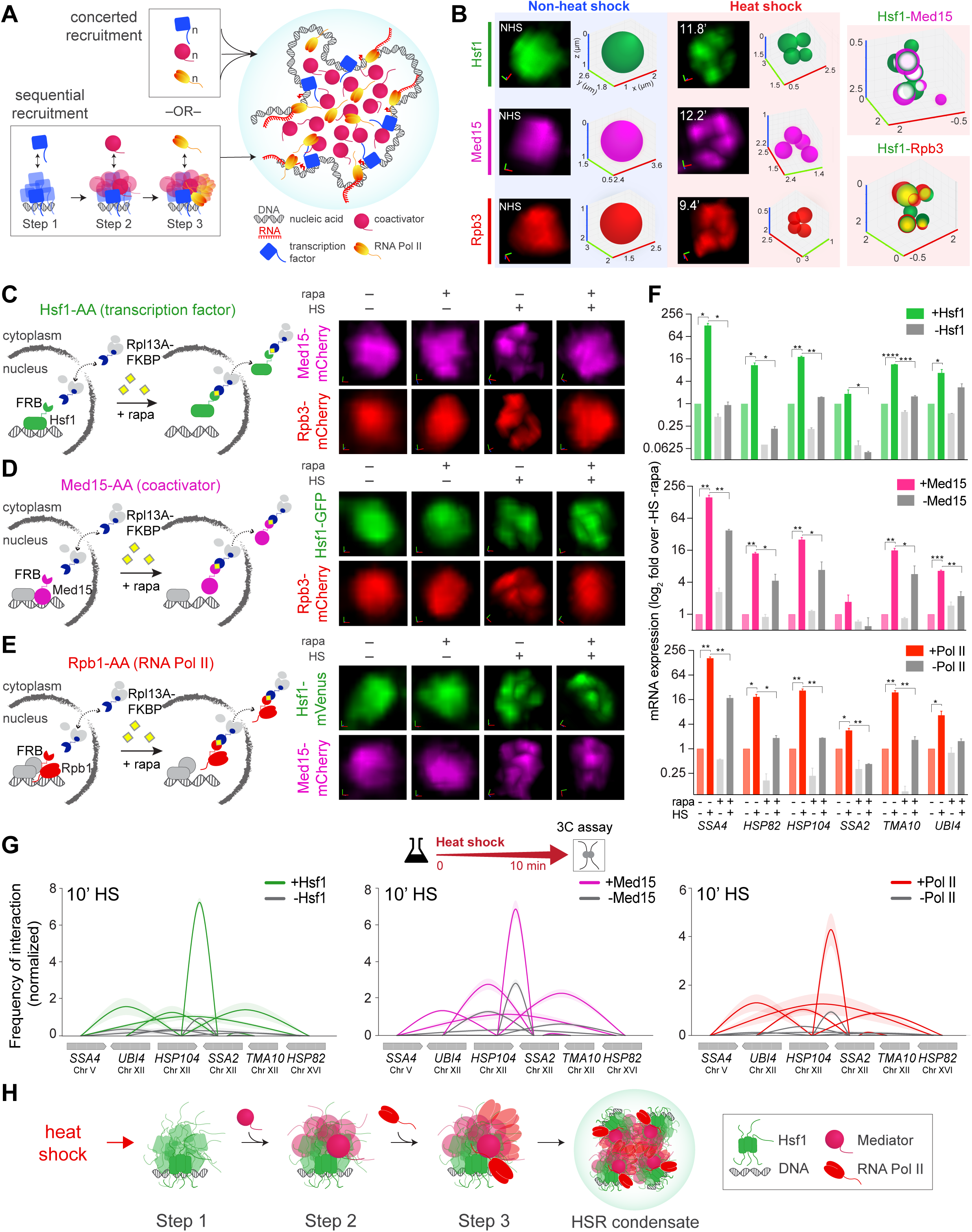
Ordered recruitment of Hsf1, Mediator, and Pol II in the HSR condensates. **(A)** Models of transcriptional condensate assembly. Concerted recruitment (top) predicts non-stochastic, simultaneous recruitment of constituent biomolecules, driving the spontaneous assembly of the condensate. The stepwise recruitment model (bottom) posits an orderly cascade of events, whereby each component is serially recruited, resulting in the gradual assembly of the condensate. **(B)** Left and middle: Representative 3D images of live yeast cells expressing either Hsf1-mVenus, Med15-mCherry, and Rpb3-mCherry before (non-heat shock) and following heat shock (39°C) for the times indicated. Shown are the zoomed-in 3D volumetric renderings of the nuclei of the three individual cells. x (red), y (green) and z (blue) axes are indicated. The Hsf1, Med15, Rpb3 nuclear foci are depicted as 3D bubble charts. Right: 3D bubble chart depiction of the colocalization of Hsf1 foci with Med15 or Rpb3 at 10 min heat shock. The colocalization data are derived from cells co-expressing Hsf1-mVenus with Med15-mCherry or Rpb3-mCherry, as shown in ^11^. **(C)** Live cell images of Med15-mCherry or Rpb3-mCherry nuclear localization in Hsf1-Anchor Away (AA) cells under control (-rapa) or Hsf1-depleted (+rapa) conditions, in presence or absence of heat shock (10 min-HS). **(D)** same as C, except Hsf1-GFP and Rpb3-mCherry nuclear localization in Med15-AA cells. **(E)** same as C, except Hsf1-mVenus and Med15-mCherry nuclear localization in Rpb1-AA cells. **(F)** mRNA expression of HSR genes in Hsf1-AA, Med15-AA, and Rpb1-AA strains heat shocked for 10 min (or not), as measured by RT-qPCR. Depicted are means + SD; n=2, qPCR=4. \*\*\*\**P* <0.0001; \*\*\**P* <0.001; \*\**P* <0.01; \**P* <0.05 (calculated using multiple unpaired t-tests). **(G)** Intergenic contacts (solid arcs) between indicated HSR gene pairs determined by TaqI-3C in 10-min HS conditions in presence or absence of Hsf1 (left), Med15 (middle), Rpb1 (right). Shown are means +/- SD (SD, arc shading); n=2, qPCR=4. **(H)** Model of stepwise recruitment of Hsf1, Mediator, and RNA Pol II in the HSR condensates. Step 1, Hsf1 independently clusters upon heat shock; step 2, Mediator clusters in the presence of Hsf1; step 3, Pol II clusters in the presence of Hsf1 and Mediator.

An alternative model of condensate assembly is the sequential recruitment model, which entails a hierarchical and ordered recruitment process, where each component condenses in a stepwise fashion (Figure 1A). The extent to which the assembly and activity of transcriptional condensates differentially depend on cooperative interactions among multiple components versus specific pairwise interactions is, however, unclear. Unravelling the process of condensate assembly, capturing the real-time dynamics, and accounting for the complexity arising from regulatory mechanisms requires investigation of condensates in living cells.

The eukaryotic heat shock response (HSR) provides an ideal experimental system for the functional analysis and molecular characterization of transcriptional condensates. First, the HSR is an evolutionarily conserved transcriptional program that controls the expression of molecular chaperones and other components of the protein homeostasis (proteostasis) network ^25–31^. Second, the HSR involves transcriptional control through condensates ^32,33^. In *S. cerevisiae* (budding yeast), Heat Shock Factor 1 (Hsf1), the master transcription activator of the HSR, forms biophysically dynamic subnuclear clusters with Mediator (Med15) and Pol II (Rpb3) (Figure 1B) ^11^. These clusters are also observed in mammalian cells ^12^. Hsf1 clusters in yeast constantly rearrange (intermix) and form at canonical Hsf1 target/HSR gene and mRNA loci, marking them as sites of active nascent mRNA production ^11^. Third, the HSR provides the opportunity to characterize the functional regulation of the condensates. Hsf1 transcriptional condensates (aka HSR condensates) involve interactions between multiple transcriptional components via their IDRs and structured domains that are susceptible to post-translational modifications ^11^. Lastly, the HSR provides the opportunity to dissect the dynamics of a physiological condensate assembly process over time. Hsf1 condensates in yeast are inducible and reversible as they rapidly form (demix) and dissolve (remix) during heat shock, with the kinetics closely paralleling HSR gene activation and attenuation ^11^. Moreover, HSR condensates promote the coalescence of chromosomally unlinked HSR genes, reorganizing the yeast 3D genome during stress ^11,34,35^.

Here, we resolve how HSR condensates assemble *in vivo.* We provide experimental evidence that HSR condensates conform to a stepwise assembly process. Using live cell imaging and transcriptomics we show that the stepwise assembly of condensates enables the graded activation of HSR gene transcription, rather than a switch-like response. We identify separation-of-function mutants of Hsf1 that fail to promote Pol II clustering yet induce HSR gene activation. Notably, the absence of Pol II condensates virtually eliminates intergenic contacts among HSR genes, and the lack of Pol II condensates and intergenic interactions is associated with a fitness cost, underscoring the unique and adaptive role of Pol II condensates in the spatial gene control that extends beyond gene activation. We propose that the serial assembly of HSR condensates enables cells to dynamically fine-tune HSR gene expression, and the emergent properties of the fully assembled HSR condensates adaptively reconfigure the 3D genome for promoting cellular fitness during stress.

## RESULTS

### Ordered assembly of HSR condensates during heat shock

To dissect how HSR condensate assembly occurs in living cells, we used the rapamycin-based “anchor away” (AA) system ^36^ to conditionally deplete Hsf1, the Mediator subunits Med15 and Ssn3/Cdk8, and the largest RNA Pol II subunit Rpb1 from individual yeast nuclei ^26,35,37,38^. We first derived strains in the Hsf1-AA background, with either Med15 or Rpb3 endogenously tagged with mCherry, then monitored Med15 or Rpb3 clustering in live cells in the presence or absence of nuclear Hsf1 (Figure 1C). Under nonstress conditions, both Med15 and Rpb3 remain diffusely localized in the nucleus, even in the absence of nuclear Hsf1. Following a brief heat shock, both Med15 and Rpb3 formed clusters, but not when Hsf1 was depleted from the nucleus, suggesting that Hsf1 is required for Mediator and Pol II condensate formation.

Next, we generated strains with Hsf1-GFP and Rpb3-mCherry in the Med15-AA background and monitored Hsf1 or Rpb3 clustering in the presence or absence of nuclear Med15 (Figure 1D). While both Hsf1 and Rpb3 formed clusters upon heat shock in the presence of nuclear Med15, only Hsf1 could cluster upon nuclear depletion of Med15, and Rpb3 remained diffuse. This suggests that heat shock-activated Hsf1 can form condensates independently of Mediator, and that Mediator is necessary for Pol II condensate formation. By contrast, in the Ssn3/Cdk8-AA strain, both Hsf1 and Rpb3 formed clusters upon heat shock in the presence and absence of nuclear Ssn3/Cdk8 (Figure S1A). Thus, the tail subunit of the Mediator complex, Med15, which has been previously implicated in directly associating with Hsf1^39^, is required for Pol II condensate formation upon heat shock, but the reversibly associated kinase module containing Ssn3/Cdk8 is dispensable, aligning with their distinct roles in HSR gene activation ^37^.

Finally, we derived Hsf1-GFP and Med15-mCherry strains in the Rpb1-AA background and monitored Hsf1 or Med15 clustering in the presence or absence of nuclear Rpb1 (Figure 1E). Both Hsf1 and Med15 formed clusters in the presence and absence of nuclear Rpb1, suggesting that Pol II is dispensable for Hsf1 and Mediator condensate assembly. Collectively, these data suggest that the assembly of HSR condensates follow an orderly cascade of events, where Hsf1 initially clusters in response to heat shock, then recruits Mediator to condense, and the Hsf1-Mediator subassembly in turn recruits Pol II to condense (Figure 1H).

Functionally, the nuclear depletion of Hsf1, Med15, and Rpb3 all reduced transcriptional induction of HSR target genes during heat shock (Figure 1F). Likewise, depletion of each of the factors reduced the frequency of both intragenic and intergenic interactions between representative Hsf1 target/HSR genes (Figures 1G, S1B), a key property of HSR gene expression measured by a highly sensitive chromosome conformation capture assay TaqI-3C ^35,40^. Together, the imaging and molecular assays in the AA strains reveal that, although the cell biological assembly of HSR condensates proceeds via discrete steps, the molecular functions of stress-induced gene activation and 3D genome reorganization require assembly of all the components.

### Hsf1 clustering upon Hsp70 release

Since Hsf1 can independently cluster, we hypothesized that the mechanisms central to regulating Hsf1 activity would also influence Hsf1 clustering. In particular, the chaperone Hsp70 directly binds to Hsf1 to repress Hsf1 activity, while phosphorylation of IDRs on Hsf1 positively regulates Hsf1 transcriptional activity ^41^.

The chaperone Hsp70 binds to Hsf1 to repress its activity ^42,43^. AlphaFold Multimer was recently shown to accurately predict interactions of structured regions with IDRs ^44,45^. As Hsf1 is known to form a trimer and Hsp70 may exist as a dimer in eukaryotes ^46–49^, we performed *de novo* modeling of Hsf1 and Hsp70 in a 3:2 stoichiometry to predict their interactions. Without specific training on complexes of Hsf1 and Hsp70, AlphaFold2 Multimer ^50^ predicted that Hsp70 interacts with Hsf1 via two distinct binding sites, the N-terminal element 1 (NE1) located in the N-terminal IDR and the conserved element 2 (CE2) in the C-terminal IDR (Figure 2A, B). This is consistent with the previous finding that mutations in either or both NE1 or CE2 short linear motifs disrupt Hsp70 binding to Hsf1. As a consequence of disrupted Hsp70 binding, Hsf1 activity is de-repressed, leading to increased basal expression of HSR genes ^42,43^.

**Figure 2.**
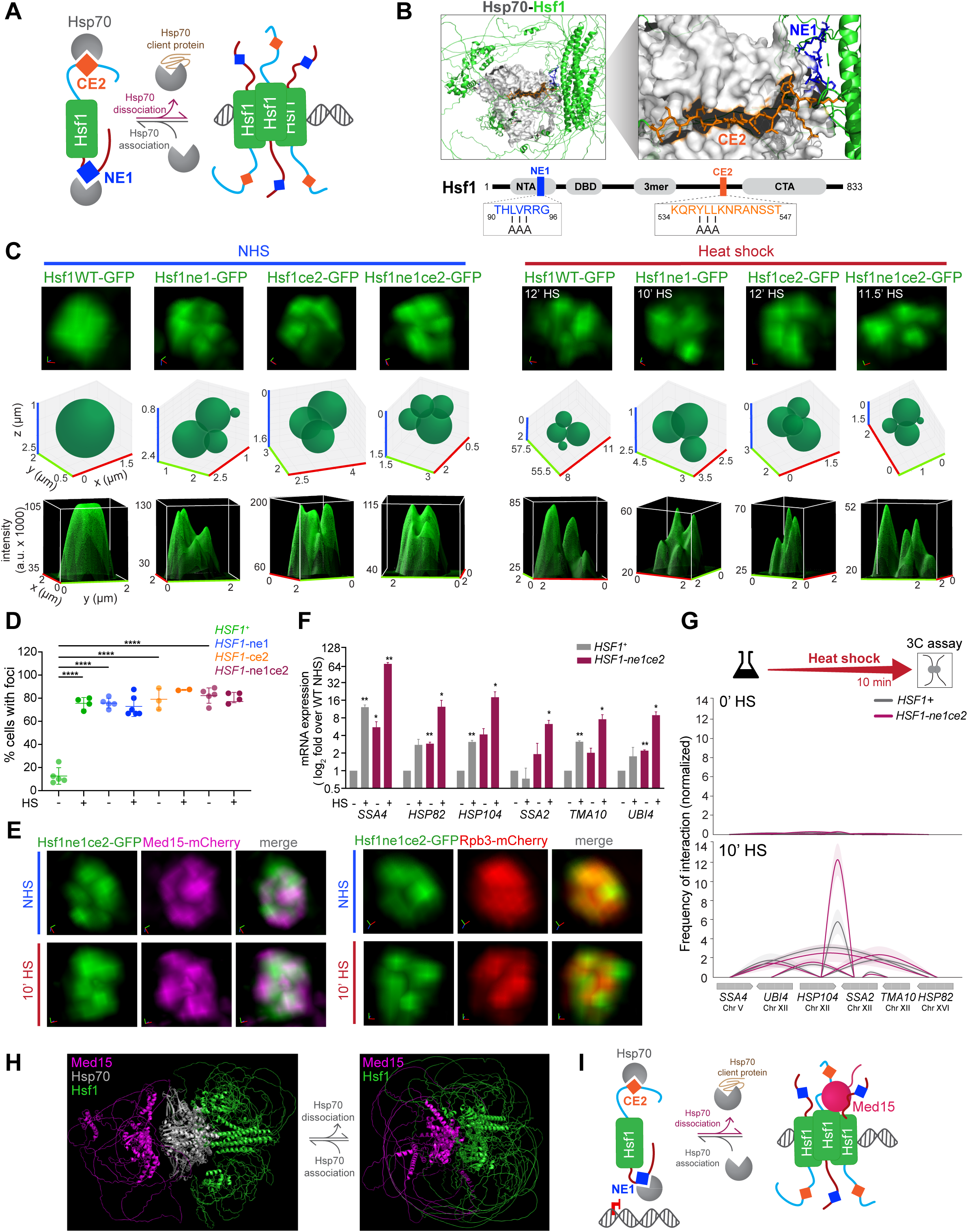
Hsf1 and Mediator cluster upon Hsp70 release. **(A)** Schematic of Hsp70 regulation of Hsf1, highlighting the N-terminal element 1 (NE1) and the conserved element 2 (CE2)—short linear motifs where Hsp70 binds to repress Hsf1. Hsf1 is activated upon sequestration of Hsp70 by its client proteins. **(B)** Top: AlphaFold2 Multimer model of the Hsf1-Hsp70 structure (3:2 stoichiometry), showing interactions between the Hsf1 NE1 and CE2 regions and Hsp70. This is not an experimentally determined structure. The predicted structures of Hsf1 (cartoon representation), Hsp70 (surface), and the NE1/CE2 sites (stick) are visualized using PyMOL; also see Data S1. Bottom: Hsf1 domain map. NTA, N-terminal activation domain; DBD, DNA binding domain; 3mer, trimerization domain; NE1, N-terminal element 1 (blue); CE2, conserved element 2 (orange); CTA, C-terminal activation domain. Boxes: NE1 and CE2 sequences and alanine substitutions in the Hsf1-ne1AAA and Hsf1-ce2AAA mutants. **(C)** Top row: 3D rendered live-cell images of Hsf1-GFP nuclear localization of wild type (WT) and Hsf1-ne1, Hsf1-ce2, and Hsf1-ne1ce2 cells under non-heat shock conditions (NHS) and heat shock (HS) conditions at times indicated. Middle row: 3D bubble charts displaying foci assignments for cells shown in the top row. Bottom row: 3D surface plots of signal intensity. **(D)** Percentage of cells with >2 Hsf1 foci per live cell quantified by FindFoci for WT and Hsf1 mutant strains under NHS and 10 min-HS conditions. Shown are means +/- SD; n= 50-65 cells/condition. \*\*\*\**P* <0.0001. *P* values were calculated by ANOVA followed by Tukey’s post hoc analysis. **(E)** Live imaging of cells co-expressing Hsf1-ne1ce2-GFP with Med15-mCherry or Rpb3-mCherry. **(F)** mRNA expression of representative HSR genes measured by RT-qPCR in WT and Hsf1-ne1ce2 double mutant in NHS and 10 min-HS conditions. Shown are means + SD; n=2, qPCR=4. \*\**P* <0.01; \**P* <0.05 (calculated using multiple unpaired t-tests). **(G)** Intergenic interactions between indicated Hsf1 target gene pairs in WT and Hsf1-ne1ce2 mutant under NHS and 10 min-HS conditions. Shown are means +/- SD; n=2, qPCR=4. **(H)** AlphaFold2 Multimer modeling of Hsf1, Hsp70, and Med15 interactions visualized in PyMOL; also see Data S2, 3. Hsp70 binding to Hsf1 is predicted to sterically occlude Med15 binding to Hsf1. **(I)** Schematic of steric hinderance by Hsp70 binding at Hsf1 to inhibit Mediator recruitment.

To determine whether binding of Hsp70 to Hsf1 regulates Hsf1 clustering and HSR condensate formation in cells, we generated mutants of Hsf1 disrupting NE1 and CE2 alone and in combination, tagged them with GFP, and expressed them from the endogenous *HSF1* promoter as the only source of Hsf1 in the cell. Live cell imaging revealed that, in contrast to the diffuse nuclear localization of wild type Hsf1, all three mutants—Hsf1-ne1, Hsf1-ce2, and Hsf1-ne1ce2—constitutively cluster into a similar number of subnuclear foci and in a similar fraction of unstressed cells, phenocopying wild type Hsf1 during heat shock. The clusters of Hsf1 mutants persisted during heat shock (Figures 2C, D, S2A). The Hsf1-ChIP experiments showed no increase in basal DNA binding at HSR genes or the non-native HSE-YFP locus (Figure S2B, C), suggesting that the constitutive clustering of Hsf1 is not associated with increased DNA binding of Hsf1 mutants. Furthermore, using the previously published RNA-seq datasets ^42^ and a single cell fluorescent reporter assay for Hsf1 activity in the mutants with individual and combined disruptions of NE1 and CE2, and leveraging the genetic interaction theory as previously applied to the HSR ^51,52^, we derived that the NE1 and CE2 sites regulate Hsf1 independently of one another (Figure S2D-F). Together, these results indicate that Hsf1 can cluster immediately following the partial release of Hsp70, and that the presence or absence of one Hsp70 binding site has no bearing on either the ability of Hsp70 to bind to the other site or Hsf1’s ability to cluster upon dissociation of Hsp70.

### Hsf1 and Mediator cluster upon complete dissociation of Hsp70

In our AA experiments, we demonstrated that Hsf1 is necessary for Med15 and Pol II condensate formation upon heat shock. To ask whether the constitutively clustered Hsf1 alone could drive the clustering of Mediator and Pol II, we derived strains with either Med15 or Rpb3 endogenously tagged with mCherry, set in the background of the Hsf1-GFP mutants with disrupted Hsp70 binding sites (Figures 2E, S2G; ^11^). While Hsf1 was constitutively clustered in all mutants, Med15 was diffuse in the Hsf1-ne1 and Hsf1-ce2 single mutants but appeared constitutively clustered in the Hsf1-ne1ce2 double mutant. In contrast, Rpb3 remained diffuse in all mutant strains in nonstress conditions. Upon heat shock, however, all the mutants displayed clustering of both Med15 and Rpb3, similar to wild type Hsf1. Thus, the microscopically similar Hsf1 clusters are in fact molecularly distinct: Hsf1 with a single mutated Hsp70 binding site clusters without Mediator or Pol II; Hsf1 with complete disruption of Hsp70 binding clusters along with Mediator but not Pol II; and the heat shock-activated Hsf1 co-condenses with both Mediator and Pol II.

While basal expression of many, if not all, HSR genes, was elevated in the Hsf1-ne1 and Hsf1-ne1ce2 mutants, each exhibited a substantial increase in inducible transcriptional activity, with the double mutant showing highest expression levels (Figures 2F, S2H). We asked if the constitutive clusters of Hsf1 mutants could drive interactions among HSR genes. Using TaqI-3C, we observed that single or double mutation of the Hsp70 binding sites did not result in any significant intergenic interactions among HSR genes. However, consistent with imaging results showing that heat shock induces full assembly of HSR condensates and high-level transcriptional activity, Hsf1 mutants showed a marked increase in both intra-and intergenic interactions among HSR genes in response to heat shock (Figures 2G, S2I, J). Thus, while the release of Hsp70 from Hsf1 and the constitutive clustering of either Hsf1 or Med15 alone are not sufficient to drive Pol II clustering or the emergence of intergenic interactions, the Hsf1 mutants retain the ability to restore wild-type functions during heat shock. This recovery is mirrored by the lack of fitness consequences of the mutants at elevated temperature (Figure S2K).

While Hsp70 is bound to Hsf1, it prevents Hsf1 clustering, which inhibits Mediator clustering. This suggests that interactions between Hsf1 and Hsp70, and between Hsf1 and Med15, are mutually exclusive. Consistent with this, AlphaFold2 Multimer modeling of Hsf1, Hsp70, and Med15 (3:2:1 stoichiometry) and Hsf1 and Med15 (3:1 stoichiometry) predicts steric occlusion of Med15 by Hsp70 (Figure 2H). Although these structural predictions with Med15 may not fully represent the functional organization in cells, which would also involve the larger Mediator subassembly, our findings suggest that interactions between Hsf1 and Hsp70 may preclude Hsf1 from engaging with Mediator under basal conditions (Figure 2I).

Taken together, our data show that Hsf1 can form clusters independently of Mediator and Pol II. The initial clustering of Hsf1 requires at least partial dissociation from its repressor, Hsp70. To partition Mediator within clusters, Hsf1 must fully unleash both its activation domains by completely dissociating from Hsp70. Hsf1-Mediator subassembly would then allow recruitment of Pol II upon heat shock, leading to full assembly of HSR condensates.

### Regulation of Hsf1 and Mediator clustering by phosphorylation

Although complete Hsp70 dissociation is sufficient to drive Hsf1 and Med15 into clusters, additional regulation is required for Pol II clustering and HSR gene coalescence. Hsf1 is phosphorylated upon activation (Figure 3A) ^53,54^, and a number of kinases have been implicated in Hsf1 regulation under different conditions across eukaryotes ^55–60^. Yeast Hsf1 can be phosphorylated on nearly 75 sites (Figure 3B), and mimicking phosphorylation is sufficient to constitutively activate Hsf1, suggesting positive regulatory potential ^41^. We tested the role of Hsf1 phosphorylation in HSR condensate formation by utilizing a phospho-null mutant of Hsf1 (Hsf1-Δpo_4_), in which alanine substitutions were made at all 152 potential phosphorylation sites, and a phospho-mimetic mutant (Hsf1-PO_4_*) with aspartate substitutions at 116 phosphorylation sites in the IDRs, as previously described (Figure 3C) ^41^.

**Figure 3.**
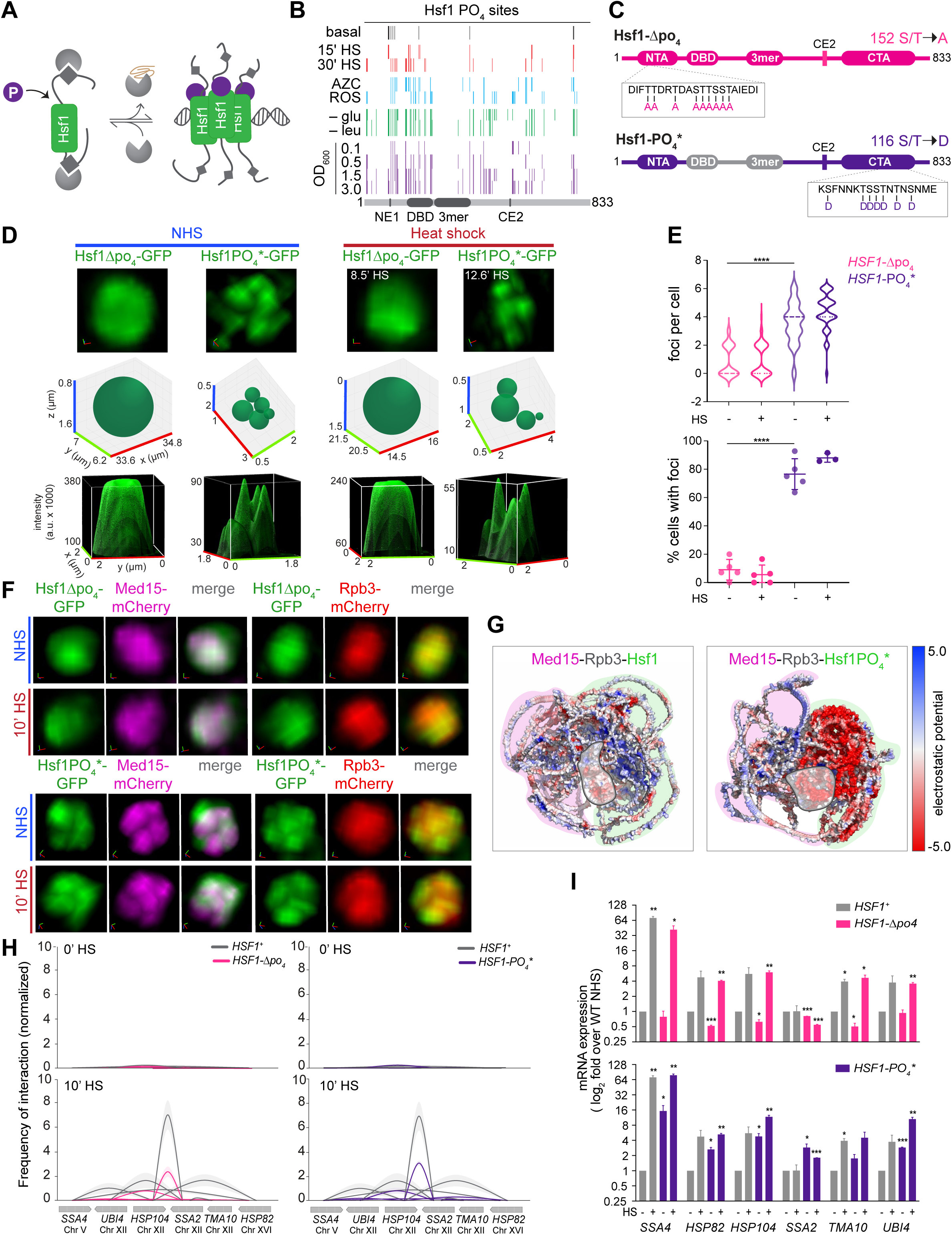
Phospho-regulation of Hsf1 and Mediator clusters. **(A)** Schematic of Hsf1 phosphorylation upon activation. **(B)** Phosphorylation sites observed in various stress and non-stress conditions shown along the primary sequence of Hsf1. HS, heat shock; AZC, Azetidine-2-carboxylic acid (1 mM); ROS, oxidative stress induced by menadione (125 µM); -glu, glucose depletion; -leu, leucine depletion. Data are derived from previously published Hsf1-3xFLAG-V5 IP/MS experiments ^41^. **(C)** Domain architecture maps of the phospho-null (Hsf1-Δpo_4_) and phospho-mimetic (Hsf1-PO_4_*) mutants. Hsf1-Δpo_4_ has all 152 possible phosphorylation sites replaced with alanine to prevent phosphorylation. Hsf1-PO_4_* has 116 phosphorylation sites in the IDRs replaced with aspartate to mimic constitutive phosphorylation. **(D)** Live imaging of Hsf1-GFP nuclear localization in Hsf1-Δpo_4_ and Hsf1-PO_4_* cells in NHS and HS conditions. Top row, 3D volumetric renderings; middle, 3D bubble charts; bottom, 3D surface plots of signal intensity. **(E)** Quantification of the number of Hsf1 nuclear foci per live cell (top) and the percentage of cells with >2 Hsf1 foci (bottom) in Hsf1-Δpo_4_ and Hsf1-PO_4_* cells under NHS and 10 min-HS conditions. Shown are means +/- SD; n=50-60 cells/condition. \*\*\*\**P* <0.0001. *P* values were calculated by ANOVA followed by Tukey’s post hoc analysis. **(F)** Top: Live imaging of cells co-expressing Hsf1-Δpo_4_-GFP with Med15-mCherry or Rpb3-mCherry under NHS and HS conditions. Bottom: same as top, except Hsf1-PO_4_*-GFP cells are shown. **(G)** AlphaFold2 Multimer models of wild type Hsf1 or Hsf1-PO_4_* in complex with Med15 and Rpb3 (3:1:1 stoichiometry) predict that phospho-mimetic residues disrupt charge block interactions of Rpb3 with Hsf1-Med15. The electrostatic surface potentials were determined using the Adaptive Poisson-Boltzmann Solver (APBS) ^89^ and visualized in PyMOL; also see Data S4, 5. **(H)** TaqI-3C analysis of intergenic contacts among HSR target genes in WT, Hsf1-Δpo_4_, and Hsf1-PO_4_* cells under NHS and HS conditions. Shown are means +/- SD; n=2, qPCR=4. **(I)** mRNA expression measured by RT-qPCR of representative HSR genes in WT, Hsf1-Δpo_4_, and Hsf1-PO_4_* cells under NHS and 10 min-HS conditions. Depicted are means + SD; n=2, qPCR=4. \*\*\**P* <0.001; \*\**P* <0.01; \**P* <0.05 (calculated using multiple unpaired t-tests).

When tagged with GFP, Hsf1-Δpo_4_ and Hsf1-PO_4_* displayed opposing nuclear distribution patterns: Hsf1-Δpo_4_ remained persistently diffuse, even upon heat shock, while Hsf1-PO_4_* was constitutively condensed, displaying a similar number of foci and fraction of cells with foci to wild type Hsf1 during heat shock (Figures 3D, E, 2D, S2A). Since Hsf1-Δpo_4_ failed to cluster, both Med15 and Rpb3 showed diffuse nuclear localization patterns despite heat shock (Figure 3F). This confirms that Hsf1 clustering is upstream of and necessary for the clustering of Mediator and Pol II. In contrast, in Hsf1-PO4* cells, Med15 was constitutively clustered along with Hsf1-PO4*, while Rpb3 remained diffuse, even upon heat shock (Figure 3F). These results indicate that phosphorylation is both necessary and sufficient for the clustering of Hsf1 and Med15, but not Pol II. Consistent with this interpretation, AlphaFold2 Multimer modeling of Hsf1 with Med15 and Rpb3 revealed compatible charge block interactions between the IDRs of wild type Hsf1 and Med15 that form a potential composite docking site for Rpb3, which is precluded by the extended acid patches in the IDRs of Hsf1-PO_4_* formed by the phospho-mimetic residues (Figure 3G). Thus, while Med15 is condensed in both Hsf1-ne1ce2 and Hsf1-PO_4_* cells, Rpb3 clusters upon heat shock in cells expressing Hsf1-ne1ce2 but not Hsf1-PO_4_* (Figures 2E and 3F). Although these structural models of isolated subunits of Mediator and Pol II with Hsf1 may not be representative of the interaction sites of the fully assembled complexes in cells, we interpret these results to suggest that phosphorylation of Hsf1 may modulate charge blocks in the IDRs that determine the affinity of Hsf1 for different surfaces on Mediator, which could potentially expose different surfaces for interaction with Pol II.

We next examined the intergenic interactions between HSR genes in Hsf1 phospho-mutants. Despite their opposing condensate phenotypes, both Hsf1-Δpo_4_ and Hsf1-PO_4_* displayed similar intergenic interaction frequencies: neither showed interactions under basal conditions, and both showed substantially reduced intergenic interactions during heat shock compared to wild type Hsf1 (Figure 3H). Surprisingly, Hsf1-PO_4_* cells showed elevated expression of key HSR genes both under basal conditions and upon heat shock (Figure 3I, bottom), despite the lack of Pol II condensates. Moreover, Hsf1-Δpo_4_ cells, also lacking Pol II condensates, induced expression of representative HSR genes during heat shock to levels comparable to wild type Hsf1 (Figure 3I, top). Thus, both Hsf1-Δpo_4_ and Hsf1-PO_4_* represent separation-of-function mutants demonstrating that Pol II condensates are dispensable for gene activation and have roles in 3D genome remodeling.

### Differential epistatic interactions impinge on condensate formation and gene activation

Since disruptions to the Hsp70 binding sites and hyper-phosphorylation of Hsf1 resulted in constitutive clustering of Hsf1 and Med15, but not Pol II, we hypothesized that combining these mutations could synergistically reprogram Hsf1 to recruit Pol II to condense in the absence of heat shock. To test this, we generated a series of combinatorial mutants of Hsf1 with disruptions to one or both Hsp70 binding sites, either in the Δpo_4_ or PO_4_* background (Figure 4A). Live cell imaging and molecular assays revealed that the phospho-mutant phenotypes were dominant to the Hsp70 binding site mutants. Similar to the Hsf1-Δpo_4_ mutant, Hsf1-ne1Δpo_4_, Hsf1-ce2Δpo_4_, and Hsf1-ne1ce2Δpo_4_ all showed diffuse nuclear localization under both basal and heat shock conditions (Figures 4B, S3A). The Hsf1-ne1ce2Δpo_4_ failed to drive HSR gene coalescence upon heat shock (Figure 4C), consistent with the absence Pol II clustering (Figure 4D). Like Hsf1-PO_4_* mutant, the Hsf1-ne1 PO_4_*, Hsf1-ce2 PO_4_*, and Hsf1-ne1ce2PO_4_* mutants clustered along with Med15 (Figure 4B, D, S3B), both in presence and absence of heat shock, and were unable to drive Pol II clustering or heat shock-dependent intra- and inter-genic interactions among HSR genes (Figures 4C, D, S3C). Thus, rather than synergistic interactions, the combined mutations displayed epistatic interactions. In terms of Hsf1 and Med15 condensate formation, mutations to the Hsf1 phosphorylation sites mask the effects of mutations to the Hsp70 binding sites.

**Figure 4.**
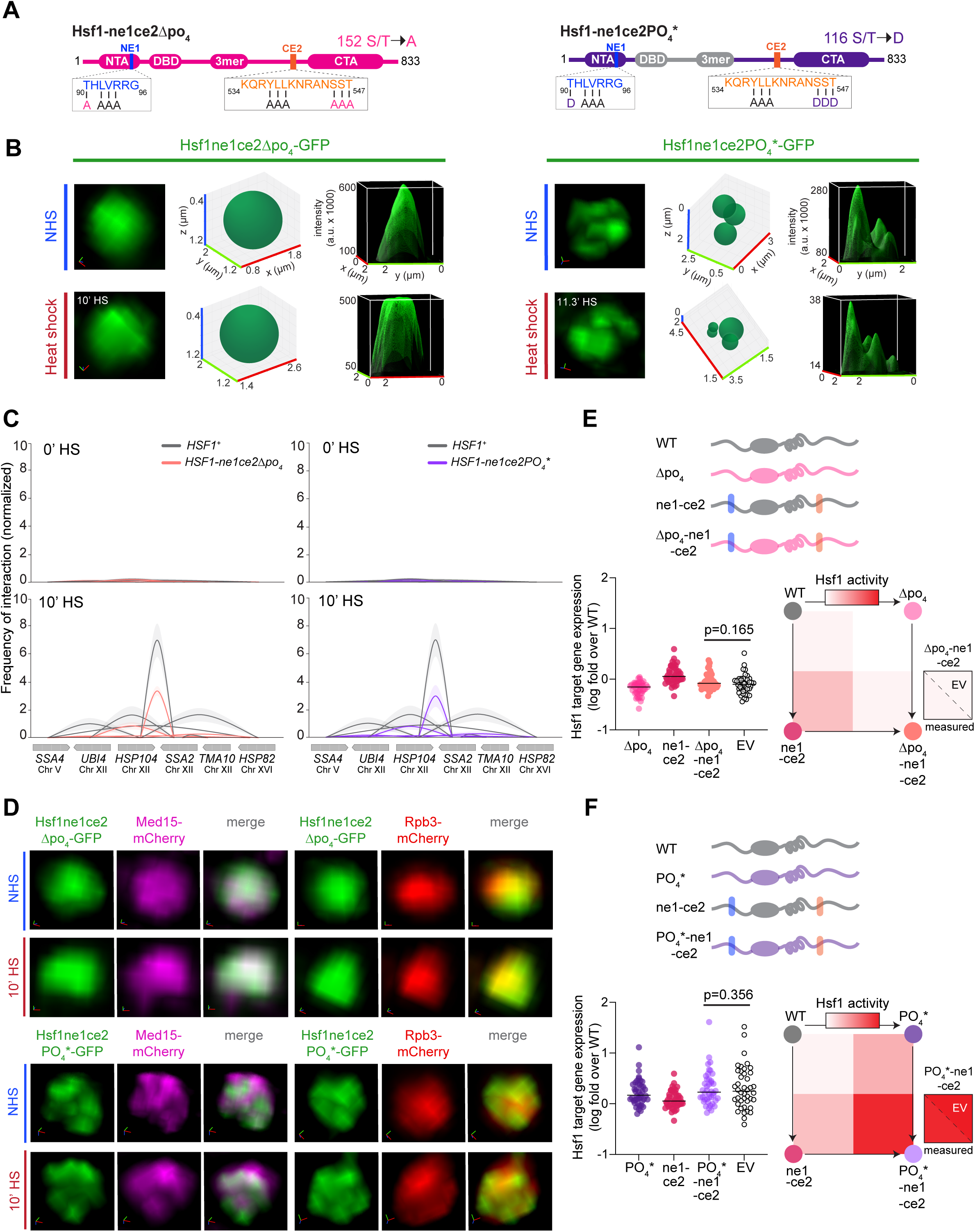
Differential epistatic control of condensate formation and transcriptional activity. **(A)** Domain architecture maps of the Hsf1-ne1ce2Δpo_4_ and Hsf1-ne1ce2PO_4_* combinatorial mutants. **(B)** Live imaging of Hsf1-GFP nuclear localization in Hsf1-ne1ce2Δpo_4_ and Hsf1-ne1ce2PO_4_* cells in NHS and HS conditions. For each mutant, left panels show 3D volumetric renderings; middle, 3D bubble charts; and right, 3D surface plots. **(C)** TaqI-3C analysis of intergenic contacts among HSR target genes in WT, Hsf1-ne1ce2Δpo_4_, and Hsf1-ne1ce2PO_4_* cells under NHS and HS conditions. Shown are means +/- SD; n=2, qPCR=4. **(D)** Top: Live imaging of cells co-expressing Hsf1-ne1ce2Δpo_4_-GFP with Med15-mCherry or Rpb3-mCherry under NHS and HS conditions. Bottom: same as top, except Hsf1-ne1ce2PO_4_*-GFP cells are shown. **(E)** Top: schematic of the double mutant cycle for Hsf1-ne1ce2 and Hsf1-Δpo_4_. Bottom left: mRNA expression of HSR genes measured by RNA-seq in the double mutant cycle for Hsf1-ne1ce2 and Hsf1-Δpo_4_. Bottom right: depiction as heat map normalized to minimal and maximal activity in the dynamic range of the assay. Shown is the comparison of the measured mRNA expression level in the double mutant with the expected value (EV) calculated from the single mutants (P value calculated using a paired t-test). For each pairwise test, n=2. **(F)** As in E, except the double mutant cycle for Hsf1-ne1ce2 and Hsf1-PO_4_* is shown.

To quantify the genetic interactions between the various Hsf1 mutations at the level of transcriptional activity, we performed double mutant cycle analyses with Hsf1-ne1ce2 mutant and the phospho-mutants ^61^. We measured mRNA expression levels of all 42 HSR genes in Hsf1-ne1ce2, Hsf1-Δpo_4_, Hsf1-PO_4_*, Hsf1-ne1ce2Δpo_4_, and Hsf1-ne1ce2PO_4_* cells, and we observed that the expression levels of the double mutants (Hsf1-ne1ce2Δpo_4_ and Hsf1-ne1ce2PO_4_*) across HSR genes were statistically indistinguishable from the expected values of the linear combination of the expression levels in the single mutants (Hsf1-ne1ce2, Hsf1-Δpo_4_, and Hsf1-PO_4_*) (Figure 4E, F). This indicates that, in contrast to condensate formation, there is no measurable epistasis between the mutations at the level of Hsf1 transcriptional activity. Thus, while the effects of mutations in the Hsf1 phosphorylation sites overpower the impact of disruptions to the Hsp70 binding sites on condensate behavior, both converge to regulate Hsf1’s transcriptional activity independently. The differential epistatic relationships at the level of condensate formation and transcriptional output provides further evidence that the process of condensate formation is distinct from gene activation.

### Graded HSR gene expression from the stepwise assembly of HSR condensates

Despite the ability to genetically separate condensate formation from transcription, the WT cells do couple the processes during heat shock. What functional role in transcriptional activation process do HSR condensates play? Our data so far indicate that the assembly of transcriptional condensates in the HSR proceeds through a series of separable steps rather than as a single concerted event. This suggests that each step in the HSR condensate assembly process may correspond to a progressive increase in HSR transcriptional activity. Indeed, RNA-seq analysis revealed a graded increase of the basal mRNA expression, spanning an order of magnitude, across the allelic series of Hsf1 mutants (Figure 5A). Principal component analysis, which supported this graded increase along PC2, clustered the mutants into groups that aligned with their condensate assembly phenotypes (Figure 5B). These results suggest that the HSR can operate as a rheostat, dynamically tuning Hsf1 activity via modulation of its propensity to assemble condensates (Figure 5C).

**Figure 5.**
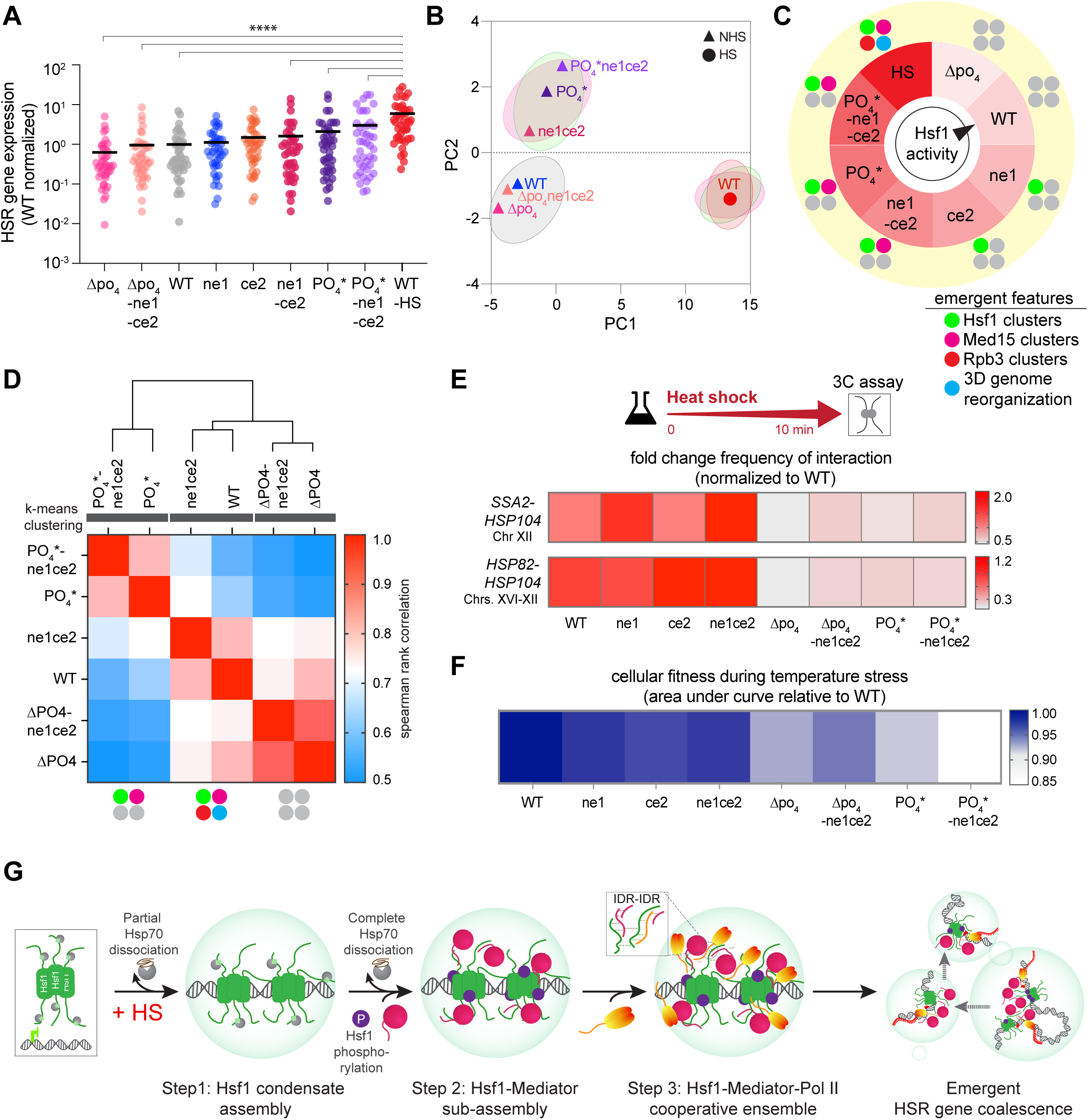
Emergent 3D genome reorganization and fitness during stress. **(A)** mRNA expression levels of HSR genes measured by RNA-seq across the allelic series of Hsf1 mutants under basal conditions compared to wild type in basal (WT) and 10 min-heat shock (WT-HS) conditions. Expression is plotted as fold change relative to the WT mean basal expression. For each pairwise test, n=2. \*\*\*\**P* <0.0001 (calculated using paired t-test). The data points for Hsf1-ne1 and Hsf1-ce2 mutants were derived from ^42^. **(B)** Principal component analysis (PCA) of RNA-seq datasets of the allelic series of Hsf1 mutants under basal conditions and of wild type in non-heat shock and 10 min-heat shock conditions. The cell-type groups are displayed as per their condensate phenotypes: green oval, cell types that can cluster Hsf1; magenta, Med15; red, Rpb3; and grey, none. **(C)** Model of Hsf1 activity as a transcriptional rheostat. **(D)** Spearman rank correlation matrix of RNA-seq datasets of wild type and Hsf1 mutants. Fold changes (10 min HS/NHS) of normalized gene counts were used to determine the optimal number of clusters grouped by k-means and hierarchical clustering. The analysis was performed with two biological replicates (n=2) for each cell type. **(E)** Fold change (10 min HS/NHS) interaction frequencies between representative chromosomally linked and unlinked HSR gene pairs for WT and allelic series of Hsf1 mutants, as measured by TaqI-3C. n=2, qPCR=4. **(F)** Integrated cell growth across the allelic series of Hsf1 mutants relative to WT during temperature stress. The quantitative growth assay was performed with two biological and six technical replicates of each cell type. **(G)** Model of stepwise condensate assembly and emergent 3D genome remodeling. Step 1 involves the initial assembly of Hsf1 condensates upon partial dissociation of Hsp70. Step 2 follows with the formation of Hsf1-Mediator subassembly upon complete Hsp70 dissociation and Hsf1 phosphorylation. Step 3 culminates in the formation of a cooperative ensemble constituting Hsf1, Mediator, and Pol II. Multiple such ensembles intermix to drive the spatial repositioning of HSR genes, promoting cellular fitness during stress.

### A specialized role of Pol II condensates in 3D genome organization and cellular fitness

The separation-of-function mutants of Hsf1 that fail to promote Pol II clustering can still induce high-level HSR gene activity, suggesting that Pol II condensates may be more involved in fine-tuning or sustaining, rather than initiating, gene activation. Instead, Pol II condensates are linked to HSR gene repositioning in response to stress. K-means clustering analysis of the heat shock-inducible mRNA expression revealed three distinct clusters that aligned with the ability of cells to form Pol II condensates. Group 1, comprising WT-Hsf1 and Hsf1-ne1ce2, can drive clustering of Pol II along with Hsf1 and Mediator upon heat shock. Group 2, which includes Hsf1-PO_4_* and Hsf1-ne1ce2PO_4_*, lacks clustering of Pol II, but not of Hsf1 and Mediator. Group 3, which constitutes Hsf1-Δpo_4_ and Hsf1-ne1ce2Δpo_4_, fails to drive the clustering of Pol II, and Hsf1 and Mediator as well (Figure 5D). Interestingly, mutants that fail to cluster Pol II (Groups 2 and 3) show a substantial reduction in intergenic interactions between HSR genes, whereas those that can cluster Pol II (Group 1) exhibit robust intergenic interactions (Figure 5E). Additionally, mutant cells lacking the ability to form Pol II condensates grow slower at elevated temperatures than WT and the mutants that can cluster Pol II (Figures 5F, S3D). These results suggest that the inability to support Pol II condensate assembly and spatial gene repositioning during heat shock is associated with reduced fitness under acute stress, underscoring an adaptive function of Pol II condensates in 3D genome reorganization during stress.

## DISCUSSION

Here, we elucidate the process underlying the assembly of transcriptional condensates in the yeast HSR. Based on our live cell investigations, we present a model in which condensates assemble sequentially. This model provides a framework for understanding the graded and tunable gene control typical of dynamic cellular environments. The prevailing view in the literature—largely derived from *in vitro* reconstitution studies—is that condensates spontaneously assemble once constituent biomolecules reach concentrations above a critical threshold. Our results indicate instead that HSR condensate assembly follows an ordered, stepwise process, wherein each key component serially condenses. This process is highly responsive to various layers of regulation, enabling precise, tunable, robust, and reversible control of HSR gene expression. In response to stress, the transcription activator Hsf1 forms clusters upon releasing the chaperone Hsp70; partial dissociation is sufficient to initiate clustering (Figure 2C). Following phosphorylation and complete dissociation from Hsp70, Hsf1 partitions the coactivator Mediator into clusters (Figures 2E, 3F), likely by unmasking interaction surfaces. The Hsf1-Mediator sub-assembly then sequentially recruits Pol II into clusters.

Building on these and the previous observations in the HSR, we propose that HSR gene regulation unfolds as follows (Figure 5G): Upon heat shock and partial release of Hsp70, Hsf1 trimers cooperatively bind to multiple Heat Shock Elements (HSEs) within the upstream activating sequences (UASs) of HSR genes ^62^, likely priming the ordered assembly of Hsf1 clusters. Phosphorylation of Hsf1 and further dissociation from Hsp70 promote non-stoichiometric—and potentially additional direct allosteric—interactions between Hsf1 and Mediator. This composite interaction surface of the Hsf1-Mediator sub-complex then facilitates the recruitment of Pol II to pre-initiation complexes at the core promoters of HSR genes. The low complexity IDRs of Hsf1, Mediator, and Pol II likely participate in multivalent interactions, helping stabilize the ordered cooperative ensemble. Many of these ensembles then come together to drive the coalescence of HSR genes across the genome.

Beyond establishing a conceptual framework for the HSR in eukaryotes, our discovery of the stepwise recruitment process offers critical new insights into broader mechanisms of gene activation. First, our findings help resolve the longstanding conflict over whether Mediator is recruited at the UASs/enhancers or the core promoters of genes, and whether Pol II helps recruit Mediator or vice versa. The cryo-EM studies of pre-initiation complexes (PICs) have captured Mediator at core promoters ^23,63^, while ChIP-seq analyses have detected substantial Mediator binding at enhancer/UAS regions, particularly for HSR genes ^11,37^. This apparent discrepancy may be explained by UAS-promoter looping, a phenomenon that HSR genes actively engage in during heat shock ^34,35^. Alternatively, Mediator may first be recruited at the UAS by the transcription activator and then facilitate the recruitment of Pol II to the core promoter. Our findings, based on conditional depletion approaches and in an allelic series of Hsf1 mutants, show that Mediator recruitment into condensates by the transcription activator Hsf1 occurs upstream of, and is necessary for, the recruitment and clustering of Pol II. Our results corroborate with and provide a cell biological manifestation of a recent single-molecule study of a synthetic gene construct, which showed that Mediator binds to the UAS-associated transcription activator independently of the core promoter, and that this binding is necessary for the subsequent recruitment of Pol II into the PIC complex ^64^ and its activation^65^.

Second, our findings provide new insights on the function of Pol II condensates in gene activation. Pol II condensates are shown to be associated with highly expressed mammalian genes and are hypothesized to be involved in transcriptional bursting, co-transcriptional splicing, and in the regulation of transcription initiation ^10,14,66,67^. Studies in live cells have shown that Pol II condensates often colocalize with Mediator subunits or transcription activators ^68^. By employing an inducible system, nuclear factor depletion strategies, and various Hsf1 mutants to study HSR condensate assembly, we were able to decouple the formation of Pol II condensates from the assemblies/co-assemblies of Mediator and transcription activators, which has allowed us to selectively parse out and focus on the specific role of Pol II condensates.

The adaptive significance of Pol II condensates appears to be linked to 3D genome reorganization. Our data suggest that Pol II condensates specifically enable intergenic interactions among HSR genes, as the mutants of Hsf1 that are incapable of driving Pol II clustering exhibit impaired 3D genome remodeling and reduced fitness under stress (Figure 5). This indicates that Pol II condensates enhance cellular adaptation, potentially by optimizing the spatial proximity and subnuclear localization of the HSR regulon. The Hsf1 target genes are shown to be targeted to the nuclear pore complexes (NPCs) ^69,70^, which may facilitate efficient export of stress-induced mRNAs, coordinating nuclear transcription with cytoplasmic translation^71,72^. Additionally, heat shock and other stresses are known to induce cytoplasmic acidification ^73,74^. These pH shifts may influence the behavior of biomolecules and promote formation of cytosolic stress granules, the adaptive condensates of mRNAs and translation initiation factors implicated in post-transcriptional gene control ^75–80^. Stress-associated pH changes may serve as a unifying signal that synchronizes transcriptional activation in the nucleus with post-transcriptional control in the cytoplasm.

Mechanistically, what drives Pol II condensate formation and HSR gene coalescence remains unclear. Beyond possible biophysical regulation, such as pH-sensitive conformational changes in Hsf1, Mediator, and/or Pol II, other factors involved in chromatin organization as well as mRNA processing and export likely contribute ^32,81^. Stress-induced transcriptional attenuation, which involves the repression of transcription factor controlling genes involved in growth and proliferation, may also help liberate Pol II to condense ^82,83^. Moreover, the mRNA transcripts emerging from the Hsf1 target genes may contribute to Pol II condensation. HSR mRNAs have been shown to form “transperons”—ribonucleoprotein particles containing mRNAs from multiple co-regulated genes ^84,85^. The rapid activation of the HSR resulting in multiplexing of mRNAs may promote Pol II condensate assembly. This would not only facilitate rapid nuclear export of the HSR mRNAs but also prevent RNA buildup, which may function as biophysical negative feedback, dispersing the condensed transcriptional machinery ^72,86^.

The ordered assembly mechanism for HSR condensates we describe enables graded control of HSR gene expression, allowing the HSR to function as a rheostat rather than a binary switch (Figure 5C). This nuanced regulation may be crucial for adaptation to varying stress intensities and durations. By contrast, 3D genome reorganization—linked to Pol II condensate assembly— acts as a switch. Under conditions when the magnitude or duration of stress exceeds a certain threshold, Pol II condensates may enable cells to maximize the adaptive capacity of the transcriptional response. The stepwise assembly mechanism we observed for HSR condensates in yeast could potentially apply to transcriptional condensates in mammalian cells, enabling graded control of transcriptional responses as the cellular signaling environment changes in development, disease, or environmental adaptation. Indeed, recent observations suggest that 3D genome organization in the context of transcriptional regulation and Pol II interaction with super-enhancer condensates are both dynamic and transient processes ^66,87^. More specifically in the context of the HSR, intergenic interactions among Hsf1 target genes and transperon formation among the target mRNAs was recently reported to be conserved in human cells, indicating that 3D genome remodeling is a conserved feature of the HSR across eukaryotes ^88^. These parallels suggest an ancient and extensive role for condensate-mediated genome organization in the eukaryotic lineage.

## METHOD DETAILS

### Yeast Strains

To tag Med15 and Rpb3 with mCherry, PCR amplicons containing the mCherry coding sequence and homology to the 3’-ends of either *MED15* or *RPB3* were amplified from the plasmid pFA6a-mCherry-hphMX6. For tagging Hsf1 with mEGFP, the PCR amplicon with homology to the 3’-end of *HSF1* was amplified from the plasmid pFA6a-mEGFP-hphMX6. These amplicons were then transformed into their respective haploid strains for in-frame genomic integration.

The plasmid pNH604-HSF1pr-HSF1-GFP was used as a template for site-directed mutagenesis to introduce NE1AAA, CE2AAA, and double mutations through QuickChange PCR. The resulting plasmids, pNH604-HSF1pr-HSF1-ne1AAA-GFP, pNH604-HSF1pr-HSF1-ce2AAA-GFP, and pNH604-HSF1pr-HSF1-ne1AAAce2AAA-GFP, were then linearized with Pme I (New England Biolabs) and transformed into the strain DPY034 for integration at the *TRP1* locus to obtain strains DPY1804, DPY1805, and DPY1806. The loss of parental HSF1 plasmid was confirmed by growth on 5-FOA media.

Plasmids pNH604-HSF1pr-HSF1Δpo4-GFP, pNH604-HSF1pr-HSF1PO4*-GFP, pNH604-HSF1pr-HSF1Δpo4ne1AAA-GFP, pNH604-HSF1pr-HSF1Δpo4ce2AAA-GFP, pNH604-HSF1pr-HSF1Δpo4ne1AAAce2AAA-GFP, pNH604-HSF1pr-HSF1PO4*ne1AAA-GFP, pNH604-HSF1pr-HSF1PO4*ce2AAA-GFP, and pNH604-HSF1pr-HSF1PO4*ne1AAAce2AAA-GFP were linearized with Fse I (New England Biolabs) and transformed in the strain DPY034 for integration at the *TRP1* locus. The loss of parental HSF1 plasmid was confirmed by growth on 5-FOA media, yielding strains SCY028, DPY1940, SCY033, SCY034, SCY035, SCY036, SCY037, and SCY038.

A complete list of strains is provided in Table S1. PCR primer sequences are provided in Table S5.

### Culture Conditions

For microscopy, cells were grown at 30°C in SDC (synthetic dextrose complete) media to early-log density (A_600_ = 0.4-0.5). For the Anchor Away experiments, rapamycin (LC Laboratories) was added at a final concentration of 1 µg/ml to early log cultures (A_600_ = 0.3-0.35) grown at 30°C in SDC. BY4742-HSF1-AA cells were incubated at 30°C for ∼90 min, yFR1324, YM114, and YM117 for ∼60 min, in presence of rapamycin before imaging ^35,37^.

For 3C, RT-qPCR, and RNA-seq analyses, cells were grown at 30°C in YPD (yeast extract-peptone-dextrose) to mid-log density (A_600_ = 0.65-0.8). A portion of the culture was maintained at 30°C as the non-heat-shocked (NHS) sample, while the remaining culture was subjected to an instantaneous 30°C to 39°C thermal upshift for the indicated duration (heat-shocked (HS) sample). For the Anchor Away 3C and RT-qPCR experiments, rapamycin was added to early log cultures (A_600_ = 0.35-0.4). Cells were grown to mid-log density at 30°C in YPD in the presence of rapamycin as described above.

For the quantitative growth assay, cells were grown to stationary phase at 30°C in YPD media. The saturated cultures were diluted to a uniform cell density of A_600_=0.15 and transferred to a 48-well microtiter plate. Cells were then grown either at 30°C for 48 h, or at 30°C for 6 h followed by a shift to 37°C for 42 h, as indicated.

For flow cytometry, cells were grown to log phase at room temperature (25°C) in SDC media. The cells were then transferred to a 96-well microtiter plate and incubated at 30°C for 1 h following the addition of cycloheximide at a final concentration of 50 µg/ml.

### Fluorescence microscopy

Fluorescence live-cell imaging was performed as previously described ^11^. Briefly, cells were grown at 30°C in SDC to early log density of A_600_ = 0.5. An aliquot of cells was immobilized on a concanavalin A-coated glass bottom dish, and fresh SDC media was added before imaging. Images were acquired on Leica TCS SP5 II STED-CW laser scanning confocal microscope (Leica Microsystems) equipped with a GaAsP hybrid detector, with STED mode disabled. Heat shock at 39**°**C was applied by heating the objective (HCX PL APOCS 63x/1.4 oil UV) with a Bioptechs objective heater system and controlling the temperature in the incubator chamber enclosing the microscope. Argon laser was used at lines 488 and 514 nm for excitation of GFP and mVenus, while mCherry was excited either at 561 nm or by using the HeNe laser at 594 nm. For the Anchor Away experiments, the SDC media in the dish was replaced with SDC supplemented with rapamycin at a final concentration of 1 µg/ml, and the cells were imaged at room temperature (∼25°C) or heat-shocked at 39**°**C for the indicated times.

All images were acquired in xyz scan mode, capturing 8 planes along the z-axis with a 0.25 µm interplanar distance. For dual-color live-cell imaging, fluorophores were sequentially scanned in two channels, with scanning modes switched on between lines to minimize bleed-through and inter-channel crosstalk. The images were deconvolved using the YacuDecu function, which applies the Richardson-Lucy algorithm to calculate theoretical Point Spread Functions (PSFs) (https://github.com/bnorthan) ^90^. The PSFs were computed based on the imaging parameters for each condition. Custom Fiji plugins were used for basic image processing, such as to colorize, split or merge channels, create composites, and adjust the brightness^91^. 3D rendering and visualization were performed using the ClearVolume plugin in Fiji ^92^.

### Chromosome Conformation Capture (3C)

TaqI-3C was performed as previously described ^11,34,35,40^. Briefly, cells were grown to mid-log density (A_600_ = 0.8) at 30°C, and either maintained at 30°C or heat-shocked at 39°C for 10 min, followed by crosslinking with formaldehyde (1% final concentration). For 3C assays involving conditional depletion by Anchor Away, cells were pre-treated with rapamycin as described above, heat-shocked at 39°C for 10 min (or not), and then crosslinked. Crosslinked cells were harvested and subjected to glass bead lysis in FA lysis buffer (50 mM HEPES pH 7.9, 140 mM NaCl, 1% Triton X-100, 0.1% Sodium deoxycholate, 1 mM EDTA, 1 mM PMSF) by vortexing at 4°C for two 20-min cycles. A 10% fraction of the chromatin lysate was digested with 200 U of Taq I (New England Biolabs) at 60°C for 7 h. Taq I was heat-inactivated at 80°C for 20 min. The digested chromatin was centrifuged, and the pellet was resuspended in 100 μl of 10 mM Tris-HCl (pH 7.5). The Taq I-digested chromatin was diluted 7-fold, and 10,000 cohesive end units of Quick T4 DNA ligase (New England Biolabs) were added for proximity ligation at 25°C for 2 h. The ligated sample was treated with DNase-free RNase (Sigma Aldrich) at 37°C for 20 min, followed by Proteinase K (Invitrogen) at 65°C for 12 h. The 3C DNA template was then extracted using phenol-chloroform and precipitated with ethanol.

The 3C interaction frequencies were quantified by qPCR on a CFX Real-Time PCR system (Bio-Rad) using Power SYBR Green PCR master mix (Fisher Scientific). Normalization controls were applied to account for: (i) variations in primer pair efficiencies; (ii) primer dimer background; (iii) differences in 3C template recovery; and (iv) ensuring a ligation-dependent 3C signal. The sequences of 3C primers used in this study are provided in Table S2. Detailed algorithms for calculating normalized 3C interaction frequencies are outlined below and in ^40^.

### Reverse Transcription-qPCR (RT-qPCR)

RT-qPCR was performed as described previously ^11^. Briefly, cells were cultured to a mid-log density (A_600_ = 0.8) at 30°C and either maintained at 30°C or heat-shocked at 39°C for 10 min. Transcription was terminated by adding 20 mM sodium azide. For RT-qPCR experiments involving conditional depletion by Anchor Away, cells were pre-treated with rapamycin, heat-shocked at 39°C for 10 min (or not), and then treated with sodium azide. Cells were harvested and lysed using glass beads in TRIzol (Invitrogen) and chloroform for 10 min at 4°C. Total RNA was precipitated with ethanol. A fraction of total RNA (∼20 µg) was treated with RNase-free DNase I (New England Biolabs) at 37°C for 15 min, followed by heat-inactivation of DNase I at 75°C for 10 min. RNA was purified using the RNA Clean and Concentrator kit (Zymo Research). 2 µg of purified RNA and random hexamers were used with the Superscript IV first-strand synthesis system (Invitrogen) for preparing cDNA.

The cDNA reaction mix was diluted 2-fold, and 2 µl of the diluted cDNA template was used for qPCR. Relative cDNA levels were quantified using the ΔΔCt method ^34^. qPCR signal from *SCR1* Pol III transcript was used as a normalization control to account for variations in cDNA recovery. The sequences of primers used for RT-qPCR analysis are provided in Table S4.

### RNA sequencing

Cells were cultured to a mid-log density (A_600_ = 0.8) at 30°C and either maintained at 30°C or heat-shocked at 39°C for 10 min. Transcription was terminated by adding 20 mM sodium azide. Cells were harvested by centrifugation, then lysed in YR Digestion buffer with Zymolyase using the YeaStar RNA kit (Zymo Research). Total RNA was isolated as per the YeaStar RNA kit manual and purified using the RNA Clean and Concentrator kit with in-column DNase I treatment (Zymo Research). 50 µg of the purified total RNA and oligo d(T)_25_-coupled paramagnetic beads were used with NEBNext High Input Poly(A) mRNA Isolation Module (New England Biolabs) to enrich for the poly(A)+ RNA. The quality, yield, and integrity of the total RNA and polyA-enriched RNA were evaluated on Bioanalyzer with the RNA 6000 Pico kit (Agilent Technologies). 100 ng of purified poly(A) mRNA was used for cDNA library preparation using the NEBNext Ultra™ II Directional RNA Library Prep kit, and libraries were barcoded using NEBNext Multiplex Oligos for Illumina (Index Primers Sets 3-4). The average fragment length, concentration, and quality of the cDNA library were analyzed with Bioanalyzer using the High Sensitivity DNA kit (Agilent Technologies). The libraries were paired-end deep sequenced on the Illumina NovaSeqX platform (NovaX-10B-300-(PE150)) at the University of Chicago Functional Genomics Core Facility (RRID:SCR_019196). For RNA-seq data analysis, see below.

### Quantitative growth assay

Cells were grown to saturation in YPD media at 30°C. The overnight grown cultures were then diluted to a density of A_600_=0.15 and transferred to a 48-well microtiter plate. The plates were incubated on the SPECTROstar Nano absorbance microplate reader (BMG LABTECH), equipped with a full spectrum UV/vis spectrometer and orbital shaking. For growth kinetic measurements in basal conditions, the plates were incubated at 30°C with shaking for 48 h, with absorbance readings taken every 30 min at 600 nm. For measurements at elevated temperature, plates were first incubated at 30°C for 6 h, then at 37°C for additional 42 h. Absorbance measurements, protocol set-up, and data acquisition were managed using the MARS data analysis software.

### Flow cytometry

For the reporter assays, cells were grown to mid-log phase in SDC media at 25°C. A 50 ul aliquot of cells was transferred into a 96-well microtiter plate containing 50 µl of SDC supplemented with cycloheximide at a final concentration of 50 µg/ml and incubated at 30°C for 1 h to allow maturation and folding of the reporter. HSE-YFP reporter activity was measured using BD Fortessa HTS 4-15 benchtop analyzer at the University of Chicago Cytometry and Antibody Technology Core Facility (RRID: SCR_017760). The data were analyzed using FlowJo.

### Chromatin Immunoprecipitation (ChIP)

250 ml culture of cells were grown to an OD_600_ of 0.8 in YPD media, heat-shocked at 39°C for 15 min (or maintained at 30°C), then crosslinked with 1% formaldehyde. Cells were filtered and then cryolysed in lysis buffer (50 mM HEPES pH 7.5, 140 mM NaCl, 1% Triton X-100, 0.1% Sodium deoxycholate, 1 mM EDTA, 2 mM PMSF, and 250 µg/ml cOmplete™, EDTA-free Protease Inhibitor Cocktail) at 4°C. The chromatin lysate was sonicated to an average fragment size of ∼250 bp using Biorupter Pico (Diagenode). A fraction of the sonicated chromatin was incubated with anti-FLAG M2 magnetic beads (Sigma Aldrich) for 16 h at 4°C. The beads were sequentially washed with lysis buffer, high salt buffer (50 mM HEPES pH 7.5, 500 mM NaCl, 1% Triton X-100, 0.1% Sodium Deoxycholate, 1 mM EDTA), and wash buffer (10 mM Tris pH 8.0, 250 mM LiCl, 0.5% NP-40, 0.5% Sodium Deoxycholate, 1 mM EDTA). Chromatin was eluted from the beads with 50 µl of 3X-FLAG peptide. RNA and proteins were removed by treatment with DNase-free RNase (final concentration of 200 µg/ml; incubation at 37°C for 1 h) and Proteinase K (final concentration of 50 µg/ml; incubation at 60°C for 16 h). The ChIP template was extracted using phenol-chloroform and precipitated in the presence of ethanol.

ChIP occupancy was quantified by qPCR. The ChIP DNA quantities were determined by interpolation from a standard curve generated with genomic DNA. The qPCR signal for each primer combination was normalized to the corresponding signal from the input DNA to account for variations in DNA template recovery.

## QUANTIFICATION AND STATISTICAL ANALYSIS

### Foci counting and characterization

Hsf1, Med15 and Rpb3 foci were detected and quantified in 3D live-cell images using the automated FindFoci plugin in ImageJ^93^, as described previously ^11^. Briefly, a region of interest was defined in each cell’s nucleus, and foci were identified across all z planes (z=8). We trained the FindFoci algorithm using the optimizer function in the plugin submenu to predict optimum parameter settings for foci assignment. The following parameters were applied to all images in this study: (a) background, one standard deviation above mean; (b) search method, above background with an optimal value of 0.3; and (c) merge option, relative above background with peak parameter of 0.2, and minimum size of 1.

We used the x, y, z position coordinates and sizes of peak intensity regions to create 3D bubble charts in MATLAB (MATLAB, 2021. *version 9.11.0.1873467* (R2021b), Natick, MA: The MathWorks, Inc.). For visualizing images as 3D surface plots, a 10 × 10 pixel square was marked around the nucleus, and representative planes for each cell were displayed as 3D surface plots using the Interactive 3D Surface Plot plugin in ImageJ (contributing author: Kai Uwe Barthel, Germany). The plot parameters for scale, rotation, smoothing, and lightning were adapted for optimal display of 3D surface plots, with consistent settings applied to both non-heat shock and heat shock conditions.

### RNA-seq analysis

Adapter sequences were trimmed from raw reads using TrimGalore v0.6.10 (https://github.com/FelixKrueger/TrimGalore) in paired-end mode (--paired) with default settings. The fasta file of *S. cerevisae* reference genome sequence (SacCer3) and its corresponding annotation gtf file were downloaded from NCBI. Genome indices were generated using STAR v2.7.11b (--runMode genomeGenerate --sjdbOverhang 100) ^94^. Trimmed FASTQ files were used to map the reads to the *S. cerevisae* reference genome with STAR v2.7.11b (--outSAMtype BAM Unsorted --alignIntronMin 10 --alignIntronMax 5000 --alignMatesGapMax 2000 --quantMode GeneCounts --genomeSAindexNbases 10), generating BAM files. These BAM files were sorted and indexed using SAMtools v1.19.2 ^95^. The raw counts of genes were extracted from the sorted BAM files using HTseq v2.0.4 (-f bam -m intersection-strict -s reverse) ^96^. The raw counts were then normalized using DEseq2 v1.34.0 ^97^.

The normalized gene counts were used for principal component analysis (PCA). k-means clustering, hierarchical clustering, and spearman rank correlation were performed using the Morpheus portal (https://software.broadinstitute.org/morpheus/). For k-means clustering, the normalized fold changes (HS/NHS) of gene counts were used to determine the optimal number of clusters through the elbow method. The hierarchical clustering dendrogram was created based on a 1-SpearmanCorrelation distance matrix.

### Quantification of 3C

The interaction frequencies determined by TaqI-3C were calculated as previously described ^11,34,35,40^. The percent digestion efficiency was calculated by amplifying a region across the Taq I restriction site using a pair of convergent primers (sequences provided in Table S3). The digestion efficiency was determined for each primer combination using the formula:

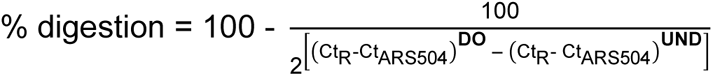

Here, Ct_R_ is the cycle threshold (Ct) quantification of the digested only (DO) or undigested (UND) templates, and Ct_ARS504_ the cycle threshold quantification of the *ARS504* locus (a region lacking a Taq I site).

For measurement of normalized frequency of intragenic or intergenic interactions, the Ct values of digested only (DO_3C_) and ligated (Lig_3C_) templates from the crosslinked chromatin, as well as the digested (DO_gDNA_) and ligated (Lig_gDNA_) genomic DNA, were used in the following formula:

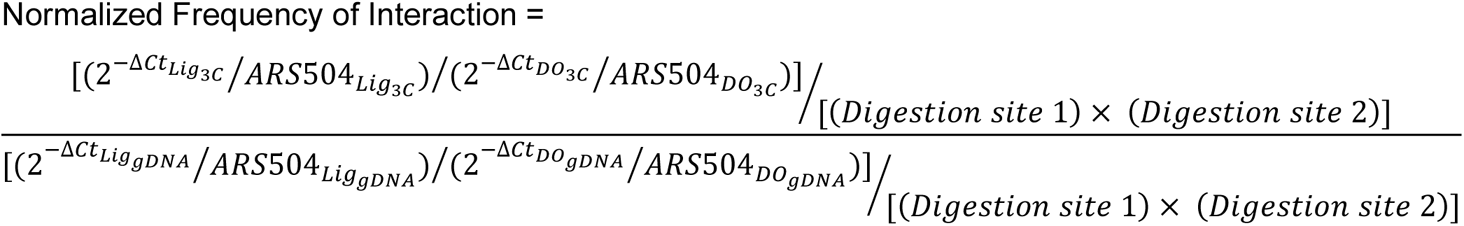

*(i) :* ΔCt values were calculated by subtracting Ct (no-template) from those of Lig_3C_, DO_3C_, or gDNA templates.
*(ii)* : 2^−Δ*Ct*^⁄*ARS*504 represents fold-over signals normalized to the *ARS504* locus.
*(iii)* : Ligation-dependent signals (LDS) are determined as the ratio of fold-over normalized signals of Lig_3C_ and DO_3C_ templates 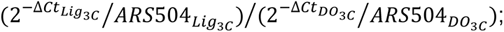 also applies to the gDNA control.
*(iv)* : LDS values were corrected for any variation in Taq I digestion efficiencies between sites 1 and 2.
*(v)* : Normalized Frequency of Interaction is the ratio of ligation-dependent signals from the 3C and gDNA control templates after correcting for differences in the digestion efficiencies.

### Quantification of genetic interactions

To quantify genetic interactions between various Hsf1 mutations, we measured Hsf1 activity using single cell fluorescent reporter and bulk RNA sequencing (RNA-seq), and we leveraged the genetic interaction theory as previously applied to the HSR ^51,52^. To this end, we assessed whether the various mutations alter Hsf1 activity independently or have interdependent effects using double mutant cycles ^61^. Based on the expression level of a reporter or target gene in wild type cells and two single mutants, we calculated the expected value of the expression level in the double mutant. If the two mutations act independently, the expected expression level in the double mutant would be the sum of the log fold changes of the individual single mutants relative to wild type. If the measured expression level diverges from the expected value, the effects of the mutations have some degree of interdependence.

### Statistical tests

Student’s *t* test (two-tailed) was used to calculate statistical significance between all pairwise comparisons (assuming parametric distributions). In Figures 2D, 3E, S2A, I, and S3C one-way ANOVA followed by Tukey’s post hoc analysis were used.

Each pairwise comparison is done with means of two independent biological samples (n=2) +SD. n.s., *P*>0.05; *, *P*<0.05; **, *P*<0.01; ***, *P*<0.001, ****, *P*<0.0001

## Supporting information

supplemental information_Chowdhary et al

## AUTHOR CONTRIBUTIONS

Conceptualization: S. C. and D. P.; Methodology: S. C., S. P., and L. D.; Formal Analysis: S. C.; Investigation: S. C., S. P., and L. D.; Resources: S. C., S. P., and L. D.; Writing – original draft: S. C. and D. P.; Writing – review and editing: S. C., S. P., L. D., and D. P.; Visualization: S. C. and D. P.; Supervision: D. P.; Funding Acquisition: D. P.

## ACKNOWLEDGEMENTS

We thank D. Gross, A. Drummond, S. Banani, A. Kainth, M. Rouches, and members of the Pincus lab for their insightful discussions and valuable input. We are grateful to A. Murugan, A. Ruthenburg, and R. Ranganathan for generously providing access to the SPECTROstar Nano absorbance microplate reader, the BioRad Real-time PCR system, and other essential lab resources. We thank M. Cianfrocco at the University of Michigan for sharing access to the COSMIC2 gateway. We greatly acknowledge the support from the University of Chicago Functional Genomics Core Facility (RRID:SCR_019196) for deep-sequencing our cDNA libraries. We are also thankful to the University of Chicago’s Integrated Light Microscopy Core (RRID: SCR_019197) and the Cytometry and Antibody Technology Core (RRID: SCR_017760) for the technical support and assistance. This work was supported by National Institutes of Health grants R01 GM138689 and RM1 GM153533 and the National Science Foundation QLCI QuBBE grant OMA-2121044 awarded to D. P.

