## supplemental information_Chowdhary et al for "Emergent 3D genome reorganization from the stepwise assembly of transcriptional condensates"

**SUPPLEMENTAL MATERIALS**

**Chowdhary et al.**

Supplemental Figures (S1-S3)

Supplemental Tables (S1-S5)

Figure S1

**A**

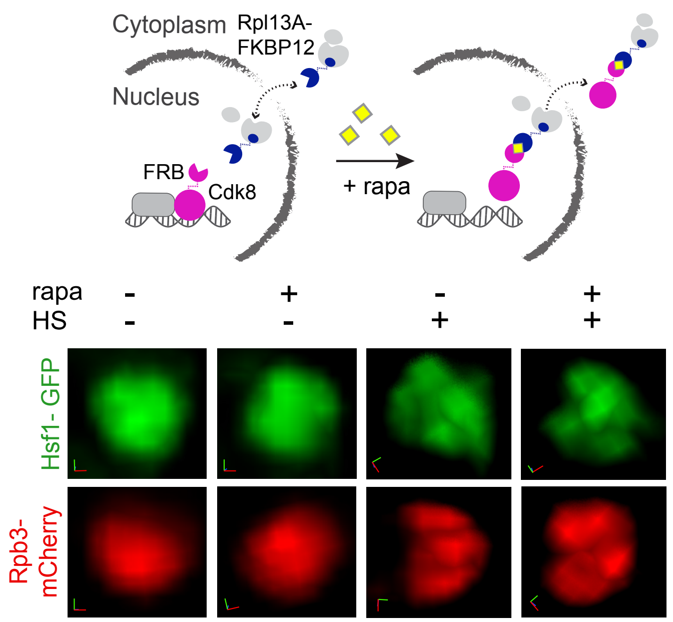

Cytoplasm
Rpl13A-
FKBP12
Nucleus
FRB
Cdk8
+ rapa
rapa - + - +
HS - - + +
Hsf1- GFP
Rpb3-
mCherry


**B**

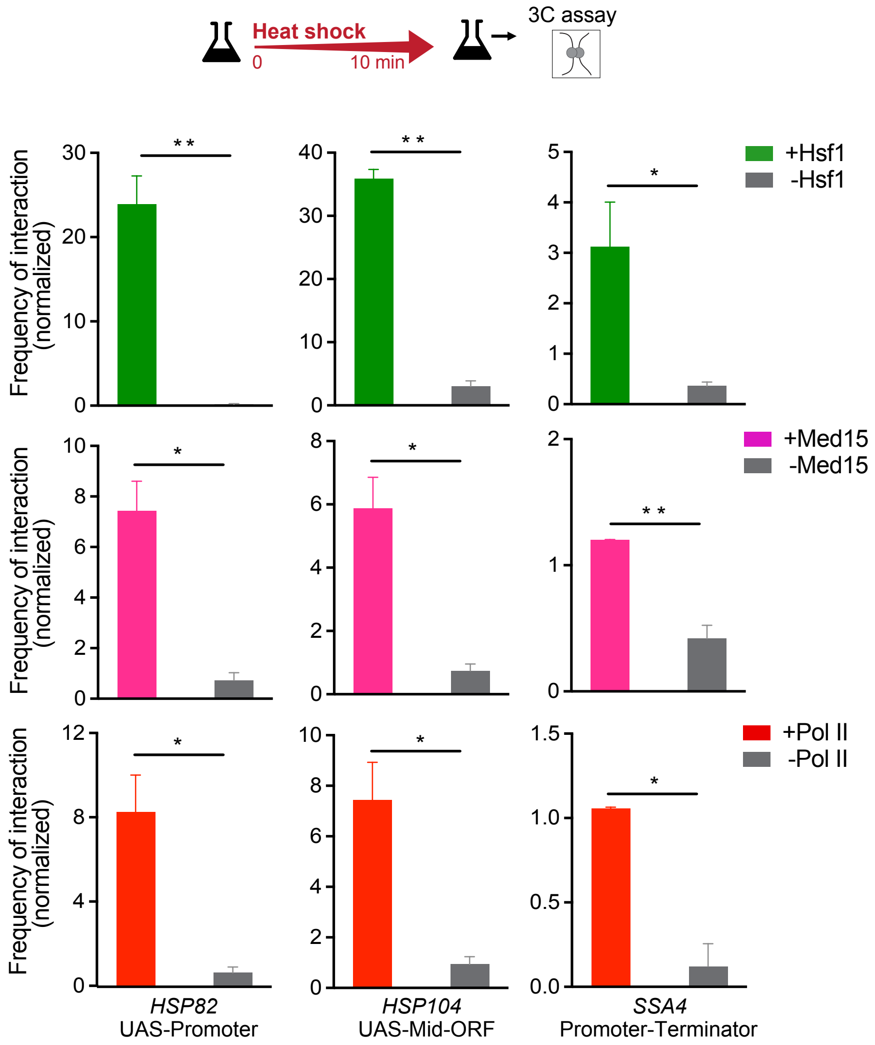

Heat shock
0
10 min
3C assay
Frequency of interaction
(normalized)
+Hsf1
-Hsf1
+Med15
-Med15
+Pol II
-Pol II
HSP82
UAS-Promoter
HSP104
UAS-Mid-ORF
SSA4
Promoter-Terminator

**Figure S1. Ssn3/Cdk8 anchor away and intragenic 3C interactions, related to Figure 1**

**(A)** Live cell images of Rpb3-mCherry (RNA Pol II subunit) or Hsf1-GFP nuclear localization in Cdk8/Ssn3-Anchor Away (AA) cells under control (-rapa) or Cdk8/Ssn3-depleted (+rapa) conditions, before and after heat shock (10 min-HS). Shown are the zoomed-in 3D volumetric renderings of the nuclei. x (red), y (green) and z (blue) axes are indicated.

**(B)** Intragenic interactions within representative Hsf1 target genes under 10 min-heat shock conditions in cells with nuclear depletion of Hsf1, Mediator (Med15), and Pol II (Rpb3), as determined by TaqI-3C. Shown are means + SD; n=2, qPCR=4. $^{**}P < 0.01$; $^{*}P < 0.05$ (calculated using unpaired t-test).

Figure S2

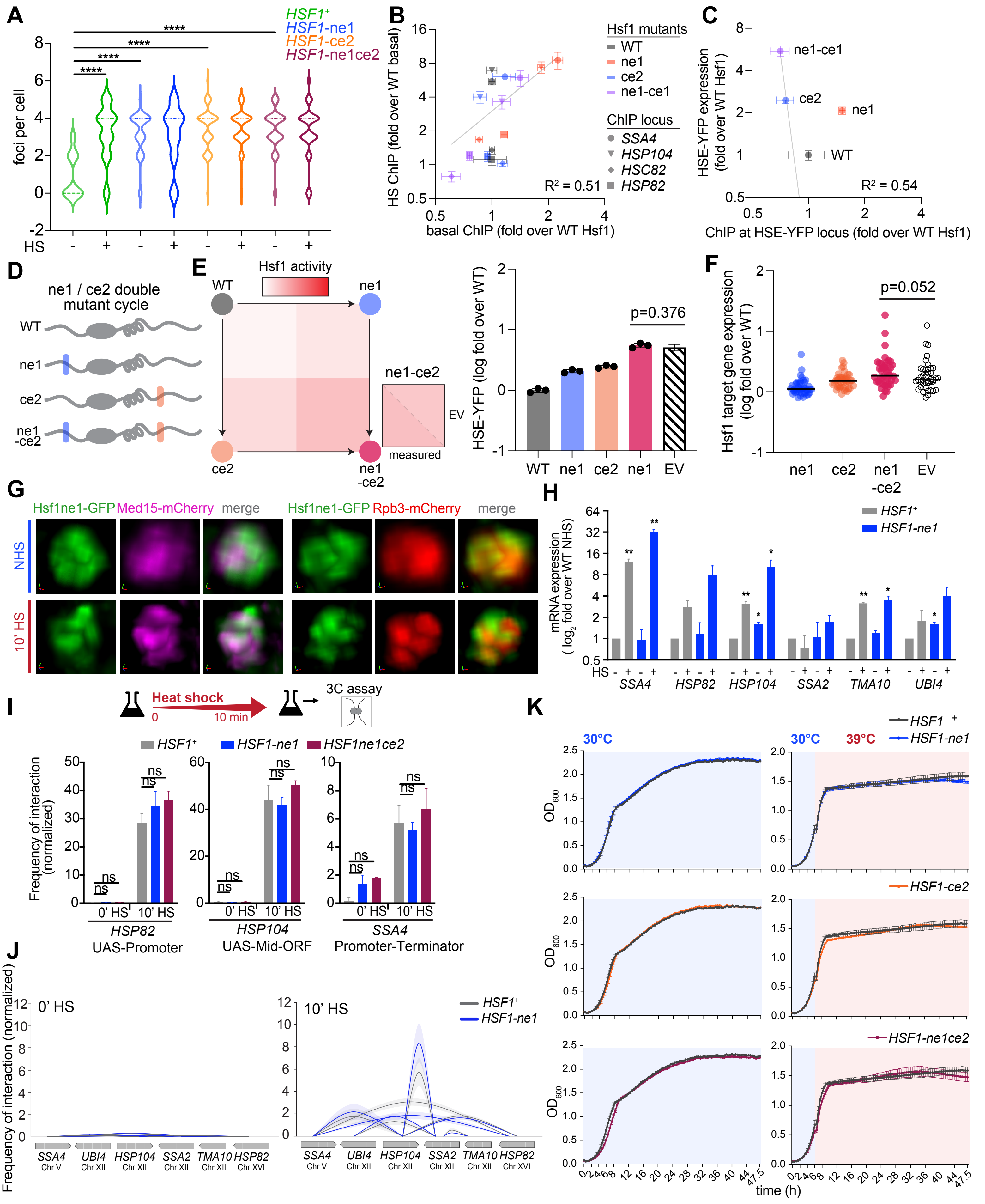
A
HSF1+
HSF1-ne1
HSF1-ce2
HSF1-ne1ce2
foci per cell
HS
B
HS ChIP (fold over WT basal)
basal ChIP (fold over WT Hsf1)
R² = 0.51
Hsf1 mutants
WT
ne1
ce2
ne1-ce1
ChIP locus
SSA4
HSP104
HSC82
HSP82
C
HSE-YFP expression (fold over WT Hsf1)
ChIP at HSE-YFP locus (fold over WT Hsf1)
R² = 0.54
D
ne1 / ce2 double mutant cycle
E
Hsf1 activity
measured
EV
HSE-YFP (log fold over WT)
p=0.376
F
Hsf1 target gene expression (log fold over WT)
p=0.052
G
Hsf1ne1-GFP
Med15-mCherry
Rpb3-mCherry
merge
NHS
10' HS
H
mRNA expression ( log2 fold over WT NHS)
SSA4
HSP82
HSP104
SSA2
TMA10
UBI4
I
Heat shock
3C assay
Frequency of interaction (normalized)
HSF1ne1ce2
0' HS
10' HS
HSP82 UAS-Promoter
HSP104 UAS-Mid-ORF
SSA4 Promoter-Terminator
J
Frequency of interaction (normalized)
0' HS
10' HS
K
30°C
39°C
OD600
time (h)

**Figure S2. Cellular, molecular, and growth phenotypes of Hsp70 binding mutants, related to Figure 2**

**(A)** Quantification of the number of Hsf1 nuclear foci in WT, Hsf1-ne1, Hsf1-ce2, and Hsf1-ne1ce2 cells under non-heat shock (NHS) and 10 min heat shock (HS) conditions. Foci were detected and quantified using the automated FindFoci algorithm in Fiji; 50-65 cells were evaluated per condition for each cell type. ****$P$ <0.0001. $P$ values were calculated by ANOVA followed by Tukey's post hoc analysis.

**(B)** Hsf1 occupancy in WT, Hsf1-ne1, Hsf1-ce2, and Hsf1-ne1ce2 cells at endogenous Hsf1 target genes. The Hsf1 ChIP signal is shown as fold change relative to WT under basal conditions versus 15 min heat shock. n=2; qPCR=6.

**(C)** Hsf1 ChIP enrichment at the HSE-YFP reporter locus. Hsf1 ChIP signals are plotted against reporter expression levels (fold change relative to WT). n=2; qPCR=6.

**(D)** Schematic of the double mutant cycle for Hsf1-ne1 and Hsf1-ce2.

**(E)** Left: HSE-YFP levels in WT, Hsf1-ne1, Hsf1-ce2, and Hsf1-ne1ce2 mutants shown as a heat map normalized to minimal and maximal Hsf1 activity in the dynamic range of the assay. Shown is the comparison between the measured HSE-YFP expression in the double mutant (Hsf1-ne1ce2) and the expected value (EV) calculated from the single mutants. Right: Quantitative analysis of HSE-YFP reporter activity in the NE1-CE2 double mutant cycle. P value was calculated using an unpaired t-test; n=3.

**(F)** Double mutant cycle analysis for NE1 and CE2 Hsp70 binding sites using published RNA-seq data (Peffer et al., 2019). $P$ value was calculated using a paired t-test.

**(G)** 3D rendered live-cell images of cells co-expressing Hsf1-ne1-GFP with Med15-mCherry or Rpb3-mCherry.

**(H)** mRNA expression of representative HSR genes measured by RT-qPCR in WT and Hsf1-ne1 mutant under NHS and 10 min-HS conditions. Shown are means + SD; n=2, qPCR=4. **$P$ <0.01; *$P$ <0.05 (calculated using multiple unpaired t-tests).

**(I)** Intragenic interactions within representative HSR genes under NHS and 10 min-HS conditions in WT, Hsf1-ne1, and Hsf1-ne1ce2 double mutant, as determined by TaqI-3C. Shown are means + SD; n=2, qPCR=4. ns (not significant), $P$ >0.05. $P$ values were calculated by ANOVA followed by Tukey's post hoc analysis.

**(J)** TaqI-3C analysis of intergenic contacts (solid arcs) among HSR genes in WT and Hsf1-ne1 cells under NHS and 10-min HS conditions. Shown are means +/- SD (SD, arc shading); n=2, qPCR=4.

**(K)** Quantitative growth curves of WT and Hsf1 mutants at 30°C and 39°C (n=2; q=6).

Figure S3

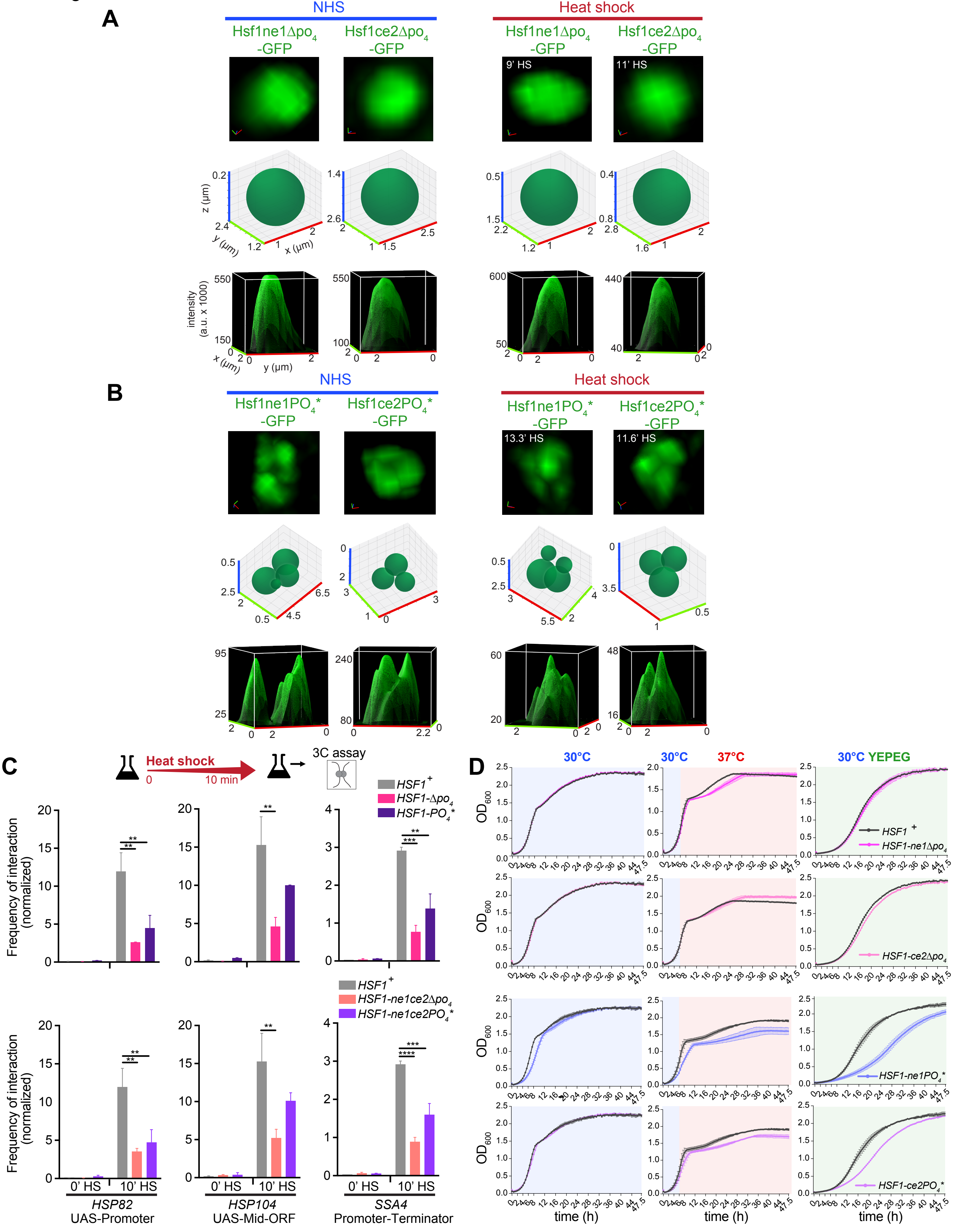
A
NHS
Heat shock
Hsf1ne1Δpo4-GFP
Hsf1ce2Δpo4-GFP
9' HS
11' HS
z (μm)
y (μm)
x (μm)
intensity (a.u. x 1000)
B
NHS
Heat shock
Hsf1ne1PO4*-GFP
Hsf1ce2PO4*-GFP
13.3' HS
11.6' HS
C
Heat shock
0
10 min
3C assay
HSF1+
HSF1-Δpo4
HSF1-PO4*
HSF1-ne1ce2Δpo4
HSF1-ne1ce2PO4*
Frequency of interaction (normalized)
0' HS
10' HS
HSP82
UAS-Promoter
HSP104
UAS-Mid-ORF
SSA4
Promoter-Terminator
D
30°C
37°C
30°C YEPEG
OD600
HSF1+
HSF1-ne1Δpo4
HSF1-ce2Δpo4
HSF1-ne1PO4*
HSF1-ce2PO4*
time (h)

**Figure S3. Cellular, molecular, and growth phenotypes of Hsf1 phospho-mutants, related to Figures 3, 4**

**(A)** Top row: Representative 3D live-cell images of Hsf1-$\Delta po_4$-GFP combinatorial mutants under NHS and HS conditions for the times indicated. Shown are the zoomed-in 3D volumetric renderings of the yeast nuclei. x (red), y (green) and z (blue) axes are indicated. Middle row: 3D bubble chart depiction of Hsf1 nuclear foci in cells shown in the top row. Bottom row: 3D surface plots displaying signal intensity in cells in top row.

**(B)** As in A, except images of Hsf1-$PO_4$*-GFP combinatorial mutants are shown.

**(C)** Intragenic interactions within representative HSR genes under NHS and 10 min-HS conditions in WT, Hsf1-$\Delta po_4$, Hsf1-PO4*, Hsf1-ne1ce2$\Delta po_4$, and Hsf1-ne1ce2PO4*, as determined by TaqI-3C. Shown are means + SD; n=2, qPCR=4. ****$P$ <0.0001; ***$P$ <0.001; **$P$ <0.01. $P$ values were calculated by ANOVA followed by Tukey's post hoc analysis.

**(D)** Growth curves of Hsf1 combinatorial mutants cultured at 30°C in glucose media, 37°C (mild stress) in glucose media, and 30°C in ethanol/glycerol (YPEG) media (moderate stress). The quantitative growth assays were performed with two biological and six technical replicates of each cell type per condition.

### SUPPLEMENTAL TABLES

#### Table S1. Yeast strains, related to STAR Methods

| Strain name | Genotype | Reference/ Source |
|---|---|---|
| DPY001 | *MATa ADE2 trp1-1 can1-100 leu2-3,112 his3-11,15 ura3-1* | Chowdhary et al., 2022 |
| DPY032 | DPY001; *HSF1-mVenus::HIS3* | Chowdhary et al., 2022 |
| SCY001 | DPY001; *MED15-mCherry::hphMX6* | Chowdhary et al., 2022 |
| SCY002 | DPY001; *RPB3-mCherry::hphMX6* | Chowdhary et al., 2022 |
| SCY003 | DPY032; *MED15-mCherry::hphMX6* | Chowdhary et al., 2022 |
| SCY004 | DPY032; *RPB3-mCherry::hphMX6* | Chowdhary et al., 2022 |
| BY4741 | *MATa his3Δ1 leu2Δ0 met15Δ0 ura3Δ0* | Chowdhary et al., 2019 |
| BY4742 | BY4741; *MATα* | Chowdhary et al., 2019 |
| BY4742-AA | *MATα tor1-1 fpr1Δ RPL13A-FKBP12::NAT MET15$^+$ lys2Δ ura3Δ his3Δ leu2Δ* | Chowdhary et al., 2019 |
| BY4742-HSF1-AA | BY4742-AA; *HSF1-FRB-yEGFP::KAN-MX* | Chowdhary et al., 2019 |
| SCY018 | BY4742-HSF1-AA; *MED15-mCherry::hphMX6* | This study |
| SCY019 | BY4742-HSF1-AA; *RPB3-mCherry::hphMX6* | This study |
| HHY212 | *MATa tor1-1 fpr1::loxP-LEU2-loxP RPL13A-2×FKBP12::loxP-TRP1-loxP ade2-1 trp1-1 can1-100 leu2-3,112 his3-11,15 ura3-1* | Anandhakumar et al., 2016 |
| YM116 | HHY212; *MED15-FRB::HIS3* | Anandhakumar et al., 2016 |
| YM117 | YM116; *LEU2* and *TRP1* excised with Cre recombinase | Anandhakumar et al., 2016 |
| SCY020 | YM117; *HSF1-mEGFP::hphMX6* | This study |
| SCY021 | YM117; *RPB3-mCherry::hphMX6* | This study |
| YM100 | HHY212; *LEU2* and *TRP1* excised with Cre recombinase | Anandhakumar et al., 2016 |
| YM114 | YM100; *CDK8-FRB::HIS3* | Anandhakumar et al., 2016 |
| SCY022 | YM114; *HSF1-mEGFP::hphMX6* | This study |
| SCY023 | YM114; *RPB3-mCherry::hphMX6* | This study |
| yFR1324 | *MATα ade2-1 trp1-1 can1-100 leu2-3, 112 his3-11,15 ura3-1 tor1-1 fpr1::NAT RPL13A-2xFKBP12::TRP1 RPB1-FRB::KAN-MX* | Chowdhary et al., 2019 |
| SCY024 | yFR1324; *HSF1-mVenus::hphMX6* | This study |
| SCY025 | yFR1324; *MED15-mCherry::hphMX6* | This study |

| | | |
|---|---|---|
| DPY182 | DPY001; *hsf1Δ::KAN HSF1pr-HSF1-GFP::TRP1* | Chowdhary et al., 2022 |
| DPY034 | DPY001; *hsf1Δ::KAN; pRS316-HSF1* | Chowdhary et al., 2022 |
| DPY1804 | DPY034; *HSF1pr-HSF1-ne1AAA-GFP::TRP1* without pRS316-HSF1 | This study |
| DPY1957 | DPY1804; *MED15-mCherry::hphMX6* | This study |
| SCY026 | DPY1804; *RPB3-mCherry::hphMX6* | This study |
| DPY1805 | DPY034; *HSF1pr-HSF1-ce2AAA-GFP::TRP1* without pRS316-HSF1 | Chowdhary et al., 2022 |
| SCY012 | DPY1805; *MED15-mCherry::hphMX6* | Chowdhary et al., 2022 |
| SCY013 | DPY1805; *RPB3-mCherry::hphMX6* | Chowdhary et al., 2022 |
| DPY1806 | DPY034; *HSF1pr-HSF1-ne1AAAce2AAA-GFP::TRP1* without pRS316-HSF1 | This study |
| DPY1959 | DPY1806; *MED15-mCherry::hphMX6* | This study |
| DPY1963 | DPY1806; *RPB3-mCherry::hphMX6* | This study |
| SCY027 | DPY034; *HSF1pr-HSF1Δpo4-GFP::TRP1* | This study |
| SCY028 | SCY027 without pRS316-HSF1 | This study |
| SCY029 | SCY028; *MED15-mCherry::hphMX6* | This study |
| SCY030 | SCY028; *RPB3-mCherry::hphMX6* | This study |
| DPY1940 | DPY034; *HSF1pr-HSF1PO4*-GFP::TRP1* without pRS316-HSF1 | This study |
| SCY031 | DPY1940; *MED15-mCherry::hphMX6* | This study |
| SCY032 | DPY1940; *RPB3-mCherry::hphMX6* | This study |
| SCY033 | DPY034; *HSF1pr-HSF1PO4*-ne1AAA-GFP::TRP1* without pRS316-HSF1 | This study |
| SCY034 | DPY034; *HSF1pr-HSF1PO4*-ce2AAA-GFP::TRP1* without pRS316-HSF1 | This study |
| SCY035 | DPY034; *HSF1pr-HSF1PO4*-ne1AAAce2AAA-GFP::TRP1* without pRS316-HSF1 | This study |
| SCY036 | DPY034; *HSF1pr-HSF1Δpo4-ne1AAA-GFP::TRP1* without pRS316-HSF1 | This study |
| SCY037 | DPY034; *HSF1pr-HSF1Δpo4-ce2AAA-GFP::TRP1* without pRS316-HSF1 | This study |
| SCY038 | DPY034; *HSF1pr-HSF1Δpo4-ne1AAAce2AAA-GFP::TRP1* without pRS316-HSF1 | This study |
| SCY039 | SCY035; *MED15-mCherry::hphMX6* | This study |
| SCY040 | SCY035; *RPB3-mCherry::hphMX6* | This study |
| SCY041 | SCY038; *MED15-mCherry::hphMX6* | This study |
| SCY042 | SCY038; *RPB3-mCherry::hphMX6* | This study |
| DPY144 | DPY001; *HSE-YFP::LEU2* | Krakowiak et al, 2018 |

| | | |
|---|---|---|
| DPY1750 | DPY144; *hsf1Δ::KAN HSF1pr-HSF1- FLAG::TRP1* | This study |
| DPY1747 | DPY1750; *HSF1pr-HSF1-ne1AAA-FLAG::TRP1* | This study |
| DPY1748 | DPY1750; *HSF1pr-HSF1-ce2AAA-FLAG::TRP1* | This study |
| DPY1749 | DPY1750; *HSF1pr-HSF1-ne1AAAce2AAA-FLAG::TRP1* | This study |

**Table S2. Forward (F) primers used for TaqI-3C, related to STAR Methods**

| Name | Sequence (5' → 3') |
|---|---|
| UBI4 F+524 | GTAAGCAGCTAGAAGATGGTAGAACC |
| HSP104 F-63 | AGGCATTGTAATCTTGCCTCAATTC |
| HSP104 F+1550 | CCCTTGATGCTGAACGTAGATATG |
| SSA2 F+198 | AGGTAACAGAACCACTCCATCTTTC |
| SSA2 F+1368 | TCTCTACTTATGCTGACAACCAACC |
| TMA10 F+811 | ATGCAAAAACACTTCCCAGAATAG |
| SSA4 F+198 | GCCTTCTTATGTGGCTTTTACTGAC |
| SSA4 F+2255 | ATAAGAAAGTCATCGCCAAACAAC |
| HSP82 F-290 | CCTCTCTCAACACAGTAATCCATAAAC |
| HSP82 F+740 | AATTAGTCGTCACCAAGGAAGTTG |
| HSP82 F+1445 | GCCAGAACACCAAAAGAACATCTAC |
| ARS504F (internal control) | GTCAGACCTGTTCCTTTAAGAGG |

**Table S3. Reverse (R) primer used for percent digestion determination in TaqI-3C, related to STAR Methods**

| Name | Sequence (5' → 3') |
|---|---|
| UBI4 R+524 | TGAATTTTCGACTTAACGTTGTCG |
| HSP104 R-63 | ATCGTTAGAGCCCTTTCTGTAAATTG |
| HSP104 R+1550 | CCACATTTTGGATCATGGAGTTG |
| SSA2 R+198 | GCTTCATATCACCTTGGACTTCTG |
| SSA2 R+1368 | TTCAATTTGTGGGACACCTCTTG |
| TMA10 R+811 | CCGGTTATAGGACCCTTATTGATG |
| SSA4 R+198 | TTTACGTCCGATCAGACGCTTAG |
| SSA4 R+2255 | GTGTTAAACTCCGGTCAAAAGAAAC |

| | |
|---|---|
| HSP82 R-290 | GAAGGACCTGGTTGGTATTAAGATG |
| HSP82 R+740 | AATGCTTAACGTACAATGGGTCTTC |
| HSP82 R+1445 | ATTCATCAATTGGGTCGGTCAAG |
| ARS504R (internal control) | CATACCCTCGGGTCAAACAC |

**Table S4. Primer used for RT-qPCR, related to STAR Methods**

| Name | Sequence (5' → 3') |
|---|---|
| HSP104 F+1922 | TTAGCTAATCCAAGGCAACCAG |
| HSP104 ORF R+1922 | ACCTTCATCGTACCCGACATAAC |
| HSP82 F+1838 | ACATGGAAAGAATCATGAAGGCTC |
| HSP82 R +1838 | AAGTCCTTGACAGTCTTGTCTTGAG |
| SSA4 F+1539 | ATCTACTGGGTAAATTTGAGTTGAGC |
| SSA4 R+1539 | CTAGCTGATTCTTAGCTTGAACACG |
| SSA2 F+1905 | CTAAATTGTACCAAGCTGGTGGTG |
| SSA2 R+1905 | AATACAGAGGAAAGCAAAAGTAAAAC |
| TMA10 F+159 | ACGAAGCAAAGTCTAACCCAAAG |
| TMA10 R+159 | TTTCATTGTTTTGGGAGTTAGAGC |
| UBI4 F+524 | GTAAGCAGCTAGAAGATGGTAGAACC |
| UBI4 R+524 | TGAATTTTCGACTTAACGTTGTCG |
| SCR1 F | CTCCACCTTCACCGCTGTTAG |
| SCR1 R | AAATATGGTTCAGGACACACTCC |

**Table S5. Primer used for construction of yeast strains, related to STAR Methods**

| **Primer Name** | **Sequence (5' → 3')** |
|---|---|
| **Primers used in construction of strains: SCY018, SCY025, DPY1957, DPY1959, SCY029, SCY031, SCY039, SCY041** | |
| Fw Med15-mCherry-hphMX6 | GTTCAGAACAATTCAATGTATGGGATTGGAATAATTGGA CAAGTGCTACTCCAGCTGAAGCTTCGTAC |
| Rv Med15-mCherry-hphMX6 | CAAACGAAGTAACTTCAAAAGTATCAAAAGTATGGAAACT TCAAATGTCCGCATAGGCCACTAGTG |
| **Primers used in construction of strains: SCY019, SCY021, SCY023, SCY026, DPY1963, SCY030, SCY032, SCY040, SCY042** | |
| Fw Rpb3-mCherry-hphMX6 | ATGCATCTCAAATGGGTAATACTGGATCAGGAGGGTATG ATAATGCTTGGCCAGCTGAAGCTTCGTAC |
| Rv Rpb3-mCherry-hphMX6 | TTCGGTTCGTTCACTTGTTTTTTTTCCTCTATTACGCCCA CTTGAGAACCGCATAGGCCACTAGTG |

| | |
|---|---|
| **Primers used in construction of strains: SCY020 and SCY022** | |
| Fw Hsf1-mEGFP-hphMX6 | AGGACCCGACAGAGTACAACGATCACCGCCTGCCCAAA CGAGCTAAGAAACCAGCTGAAGCTTCGTAC |
| Rv Hsf1-mEGFP-hphMX6 | ACTATATTAAATGATTATATACGCTATTTAATGACCTTGCC CTGTGTACCGCATAGGCCACTAGTG |
| **Primers used in construction of strains: SCY036 and SCY038** | |
| Fw Hsf1∆po4-ne1AAA-GFP_QC | TCATGATGCTAGAAAACCAGCTGCTGCTGCTCATGCAG CAGCAAGAG GTGCTCCATTGGGTATTTATCAAGC |
| Rv Hsf1∆po4-ne1AAA-GFP_QC | GAGCAGCAGCAGCTGGTTTTCTAGCATCATGAA |
| **Primers used in construction of strain SCY037** | |
| Fw Hsf1∆po4-ce2AAA-GFP_QC | GGTGCACCTATAAGACCGTACAAGCAAAGGGCTGCTGC TAAGAATCGTGCAA ACGCGGCTGCAGCTGCAGAG |
| Rv Hsf1∆po4-ce2AAA-GFP_QC | CTTGTACGGTCTTATAGGTGCACCAGCATTGTTATTCG |
